# Host immunosenescence compromises *Mycobacterium tuberculosis* clearance

**DOI:** 10.1101/2023.02.20.529217

**Authors:** Falak Pahwa, Shweta Chaudhary, Ashish Gupta, Shivam Chaturvedi, Ranjan Kumar Nanda

## Abstract

Immunosenescence increases susceptibility to infectious diseases like tuberculosis (TB) in older subjects (≥60 years) and may impact containment of *Mycobacterium tuberculosis* (Mtb) during therapeutic intervention. A deeper understanding of cellular and molecular changes with age could inform new strategies to improve therapeutic outcomes. Here, we monitored the immunopathology, frequency and functionality of immune cells across extreme age groups of C57BL/6 mice following low aerosol dose infection (100-120 cfu) with Mtb H37Rv and treatment with rifampicin and isoniazid (RIF-INH). Up to 6 weeks of infection, tissue (lung, spleen and liver) mycobacterial load in old (17-19 months; M) and aged (31M) C57BL/6 mice was similar compared to young (2-4M) mice. However, at two weeks post-treatment, older mice showed a slower rate of Mtb clearance in the lungs. Old Mtb-infected mice had higher splenic T-follicular cytotoxic (T_FC_)-like cells and proteomic analysis of flow-sorted CD4^+^CD44^+^ T cells revealed deregulated mitochondrial proteins (4-hydroxy-2-oxoglutarate aldolase, aspartate aminotransferase and prostaglandin E synthase), pointing to impaired mitochondrial function. Collectively, these findings suggest that age-associated immune alterations may impair immunometabolic processes contributing to delayed Mtb clearance. These results highlight the potential importance of targeting immunometabolic dysfunction to improve TB treatment outcomes in older populations and reduce morbidity.

**Summary:** The elderly population is particularly susceptible to tuberculosis (TB), making it crucial to understand the cellular and molecular mechanisms contributing to the decline in immune responses with age. Evaluating the immunopathology, frequency and functionality of immune cells and Mtb-specific antibody responses across different age groups is essential for developing adjunct therapies for geriatric TB patients.

## Introduction

A recent WHO report predicts that, by 2030, one in six people globally will be over 60 years of age^1^. The elderly population is particularly vulnerable to infectious diseases such as tuberculosis (TB) due to age-associated decline in immune function, often referred to as immunosenescence^2^. *Mycobacterium tuberculosis* (Mtb), the causative pathogen of TB, also triggers hypermethylation and transcriptional alterations in host immune cells, affecting the immune system^3^. Older adults not only exhibit reduced vaccine effectiveness but also show higher susceptibility to TB, with epidemiological studies indicating 2–3 times higher TB incidence and up to four times greater mortality than younger TB patients. In several East Asian regions, including Japan, Korea, and Hong Kong, 45–69% of TB cases were reported in 2020-21 from older age groups^4^. Similarly, projections from Taiwan calculated that 78% of new TB cases by 2035 will be from the geriatric population, which currently contributes to 82.1% of TB-associated mortality^5,6^. These findings underscore the growing clinical relevance of TB in an aging world.

Despite this growing burden, current TB research relies largely on young adult mouse models (6–8 weeks old, equivalent to ∼18-year-old humans), which do not adequately capture the immune landscape of older hosts^7^. Evidence from studies suggests that old (>18 months; M) mice exhibit higher bacterial burden, delayed CD4^+^ T cell responses, and altered immune activation compared to younger counterparts^8,9,10^. However, focused studies addressing how immunosenescence shapes bacterial clearance, immune cell dynamics, and host responses during TB treatment remain scarce. To address this gap, we examined the impact of aging on TB infection and treatment outcomes using a murine model. Young (2–4M), mid-aged (9-12M; 38-44 human years), old (17–19M; ∼54–60 human years), and aged (31M; >80 human years) C57BL/6 mice were infected with a low aerosol dose of Mtb H37Rv. Following treatment with rifampicin and isoniazid (RIF-INH), we monitored bacterial clearance kinetics across tissues and profiled systemic and tissue-specific immune responses, including circulating cytokines, Mtb-specific antibodies, liver micronutrients, and the proteome of splenic CD4^+^CD44^+^ T cells. Through this approach, we aim to identify age-associated processes that compromise treatment efficacy, which can inform strategies to reduce relapse risk in geriatric TB patients.

## Results

### Lung Mtb burden in C57BL/6 mice is unaffected by age or sex till 6 weeks post-infection

The influence of host age and sex on susceptibility to Mtb infection remains an area of interest, particularly at early time points. Thus, female and male C57BL/6 mice of wider age groups were aerosol infected with low-dose of H37Rv (Fig. 1A). Mice were categorized as young (2-4M), middle-aged (9M females/12M males) and old (17-19M males). Prior to Mtb infection, body weights were higher in the middle-aged and old mice than in younger controls (Supplementary Fig. S1A). During the study period, old mice lost ∼3% weight, whereas young mice gained ∼8% weight (Fig. 1B). Young female mice gained more body weight (∼20%) than males (∼10%, Fig. 1B). At 6 weeks post infection (w.p.i.), middle-aged and old male mice showed significant body weight loss (Fig. 1B). Our findings reveal that, despite age-related variations in body weight, lung Mtb burden progressively increased up to 4 w.p.i. before plateauing, with no significant differences attributable to age or sex by 6 w.p.i. (Fig. 1C).

**Figure 1.**
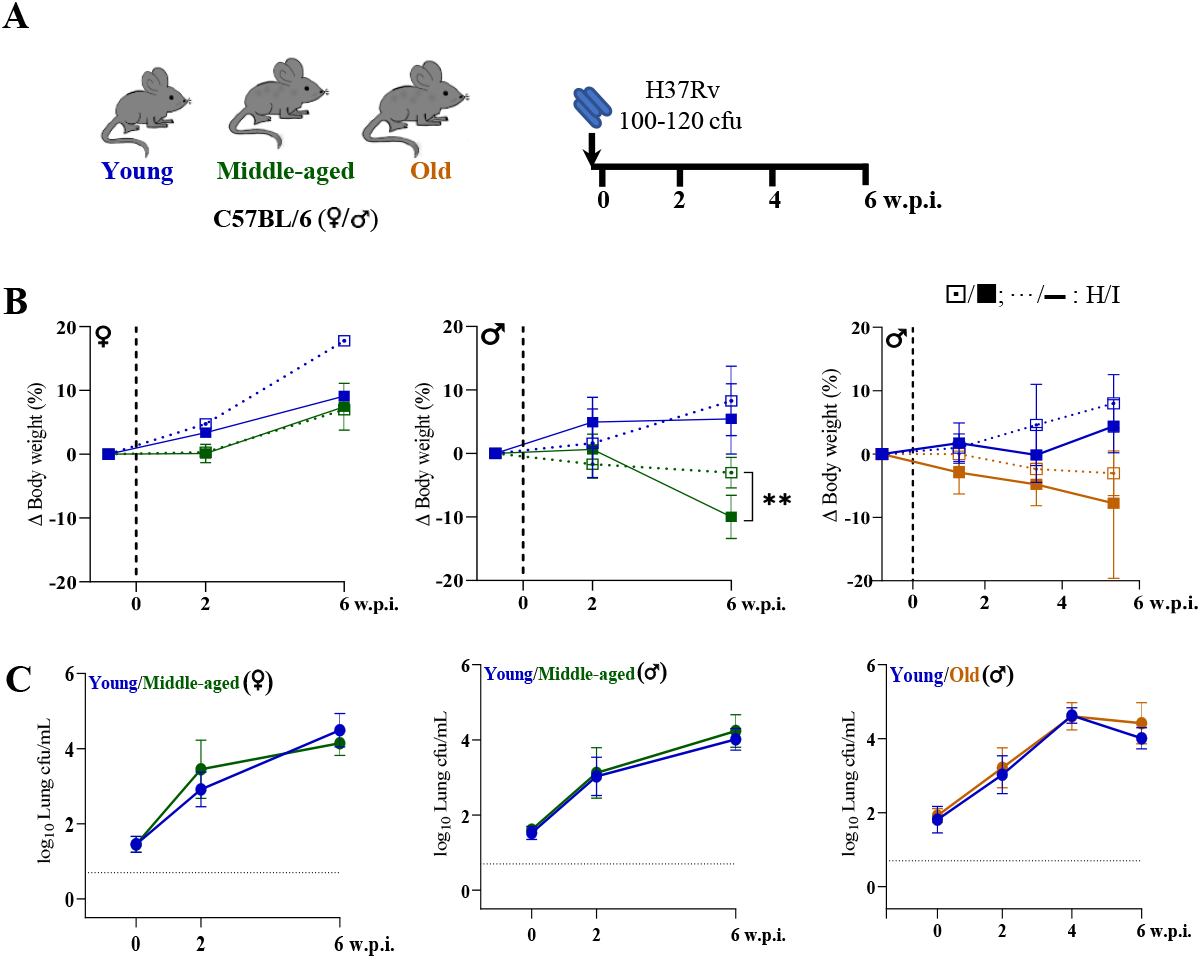
Lung Mtb burden remained similar till 6 weeks post-infection across age groups and sexes in C57BL/6 mice. **A**. Schematic of the experimental design used for Mtb H37Rv infection of C57BL/6 mice [female (**♀**) and male (**♂**)] of different age groups. **B**. Changes in body weight of mice groups (healthy: H, …. ; infected: I, —) are presented, n=5-10/age group/condition; dashed vertical line represents day of infection. **C**. Lung bacterial burden (in log_10_cfu/mL) in mice post Mtb infection; n=5-10/timepoint/age group/condition; dashed horizontal line represents limit of detection (LOD). Young (2-4 months) in blue, middle-aged (9-12 months) in green and old (17-19 months) mice in brown; cfu= colony forming unit; w.p.i.= weeks post infection; p-value: ** <0.005 at 95% confidence interval by Mann-Whitney test. Data is represented as mean ± SD. *Additional details available in Supplementary figures S1 and S2* .

### Age-dependent delays in TB treatment responses are associated with altered liver function, immune dysregulation and inflammaging

To evaluate the impact of age on treatment response, a subset of mice received RIF-INH following Mtb infection. Remarkably, old mice retained significantly higher (∼3 log_10_ cfu) lung Mtb load at 2 w.p.t. compared to the young group (∼1.5 log_10_ cfu). And by 4 weeks, both mice age groups achieved complete bacterial clearance, i.e. resulting below the limit of detection (LOD; Fig. 2A and 2B). Multiple foci of inflammatory cells and granulomatous lung lesions were observed with prominent foamy cells later (at 6 w.p.i.) in old mice but earlier (by 4 w.p.i.) in young controls (Supplementary Fig. S1). Lung inflammation was resolved in young mice by 4 w.p.t. but persisted in old mice (Supplementary Fig. S1). Also, middle-aged mice were able to resolve the inflammation by 4 w.p.t. irrespective of sex (Supplementary Fig. S2). In extrapulmonary sites, like spleen and liver, mycobacterial clearance was observed by 2 w.p.t. in both the age groups (Fig. 2B).

**Figure 2.**
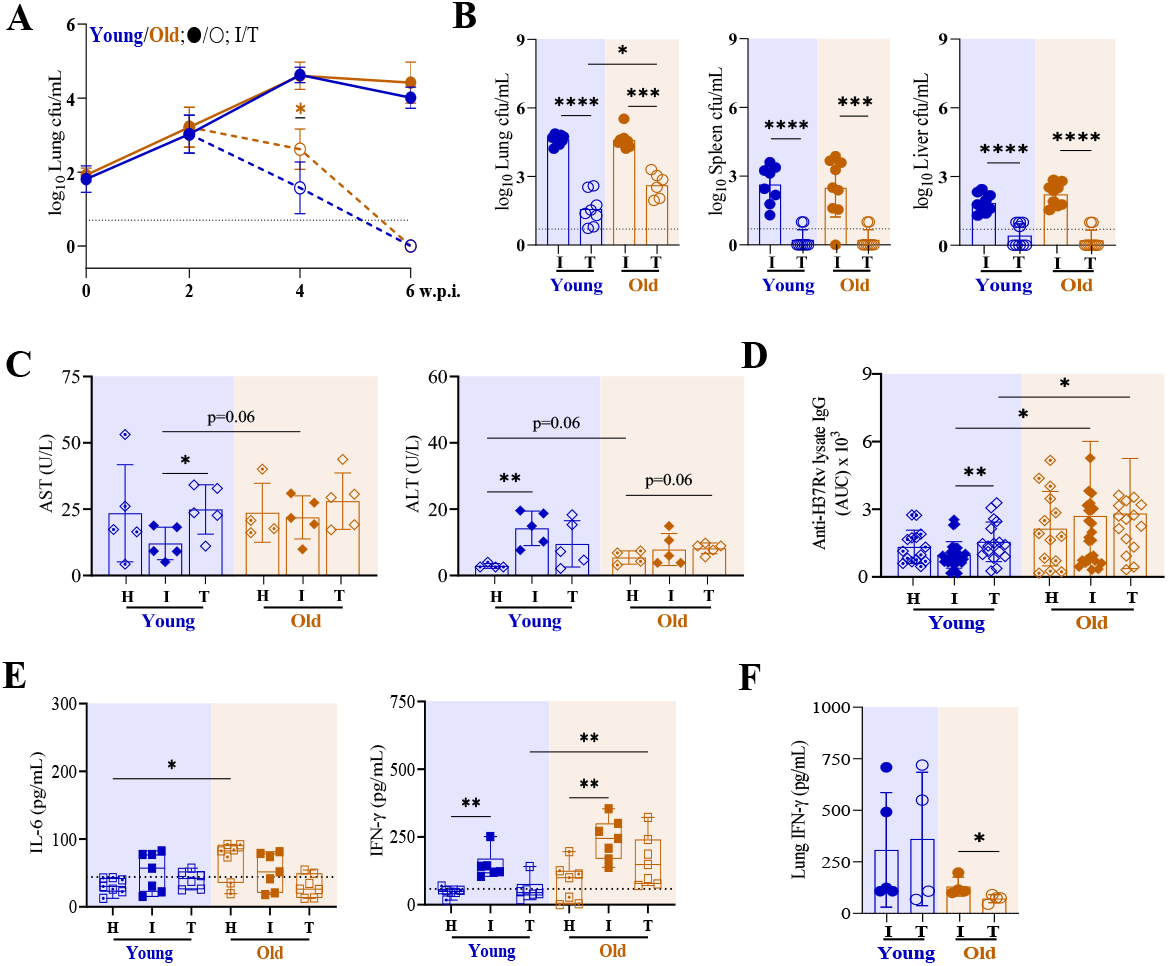
Old C57BL/6 mice receiving two weeks of RIF-INH treatment showed delayed lung Mtb clearance. **A**. Lung bacterial burden (in log_10_cfu/mL) in male C57BL/6 mice belonging to infected (I, at 2, 4 and 6 w.p.i.) and treated (T, at 2 and 4 w.p.t.; treatment started at 2 w.p.i.) groups; n=10/timepoint/age group/condition; dashed horizontal line represents limit of detection (LOD). A set of two independent experiments is depicted. **B**. Bacterial load (in log_10_cfu/mL) in the lung, spleen and liver; individual data points at 4 w.p.i./2 w.p.t. presented in histograms (n=10/age group/condition). **C**. Serum Alanine transaminase (ALT) and Aspartate aminotransferase (AST) activity in mice of different groups (n=5/age group/condition)) at 4 w.p.i./2 w.p.t.. **D**. Area under the curve (AUC) showing Mtb-specific (H37Rv lysate) IgG quantification in serum of healthy: H (n=15/age group), I (n=27-28/age group) and T (n= 16-17/age group) at 4 w.p.i./2 w.p.t.. **E**. Serum cytokines (in picogram per milliliter): IL-6, IFN-γ in the different mice groups (n=6-7/age group/condition) at 4 w.p.i./2 w.p.t.. **F**. Lung IFN-γ in the different mice groups (n=4-5/age group/condition) at 4 w.p.i./2 w.p.t.. Young (2-4 months) in blue and old (17-19 months) mice in brown; cfu= colony forming unit; w.p.i.= weeks post infection; w.p.t.= weeks post treatment; p-values: * ≤0.05, ** <0.005, *** <0.0005 and **** <0.0001 at 95% confidence interval by Mann-Whitney test. Data is represented as mean ± SD. *Additional details available in Supplementary figures S1 and S4*.

Liver enzymes play a critical role in drug metabolism, and metal ions play a significant role in their function. Age-related changes may influence this variability in liver function and micronutrient status^11^. Thus, we monitored the liver micronutrient levels of these mice groups using inductively coupled plasma mass spectrometry (ICP-MS, Supplementary Fig. S3). Old mice had higher hepatic Cu (4.28×), low Zn (0.14×) levels with similar Fe and Se levels compared to controls (Supplementary Fig. S3). Also, old mice receiving treatment showed higher hepatic Cu levels (Supplementary Fig. S3). Liver micronutrient status impacts activities of hepatic enzymes like alanine aminotransferase (ALT) and aspartate aminotransferase (AST), commonly used markers of liver function and injury. Old mice exhibited higher serum ALT and AST activities than young mice (Fig. 2C). Mtb infection did not impact serum AST or ALT activity in old mice, but ALT activity was higher in younger mice post-infection (Fig. 2C). Strikingly, old mice receiving treatment had higher serum AST activity (Fig. 2C).

To assess age-related differences in humoral immunity against Mtb, serum anti-H37Rv specific IgG titers were measured. The overall IgG response was significantly higher in old mice compared to young mice (Fig. 2D). Importantly, old mice showed higher IgG responses at a later time point i.e. 4 w.p.t., while young mice showed it at 2 w.p.t. (Fig. 2D and Supplementary Fig. S4). While robust humoral responses are beneficial for pathogen control, the delayed yet heightened antibody response in old mice could indicate persistent antigenic stimulation and likely reflects altered immune regulation with aging. Circulatory IL-6 levels were higher in old mice, consistent with a state of chronic low-grade inflammation or inflammaging (Fig. 2E). Higher circulatory IFN**-γ** levels were observed in Mtb-infected young and old mice, which resolved partially in treated groups (Fig. 2E). We observed higher lung IFN-γ levels in Mtb-infected young mice compared to old mice (Fig. 2F). Old mice receiving treatment had declined lung IFN-γ levels (Fig. 2F). Thus, the interplay between micronutrient imbalance, delayed humoral immunity and inflammaging, likely underpins the compromised mycobacterial clearance observed in older hosts.

### Age-related T cell dysregulation impairs immune control of Mtb infection

CD4^+^ T cells are critical for Mtb control; however, in older hosts, accumulation of dysfunctional T cells leads to a compromised immune response. T cells were profiled using activation markers (CD44, PD1), homing receptor (CXCR5), senescence (KLRG1, CD153) and lineage-defining transcription factors (Tbet for T_H_1 and FoxP3 for T_reg_ cells) and a significant change in their distribution was observed in old mice (Supplementary Fig. S5). Notably, high expression of CD44, CXCR5, PD1 and CD153 in CD4^+^ and CD8^+^ T cells of old mice was observed (Supplementary Figs. S5 and S6). Increased circulatory CD4^+^KLRG1^+^PD1^+^ cells were observed in old mice, indicative of exhausted T cell phenotypes (Supplementary Fig. S5).

Next, we monitored the immune cell distribution in lungs and spleen of Mtb-infected and RIF-INH treated mice groups (Fig. 3A, 3B and Supplementary Fig. S7). Old mice showed a lower proportion of splenic CD4^+^, higher CD8^+^ T cells, along with higher CXCR5 and PD1 expression compared to young mice (Fig. 3C and Supplementary Fig. S7). Interestingly, lungs of old mice showed accumulation of CD4^+^CD44^+^CXCR5^+^ (T_FH_)-like cells, similar to young mice post Mtb infection (Supplementary Fig. S8). Also, CD8^+^CD44^+^CXCR5^+^ (T_FC_)-like cells were observed to be higher in the lungs and spleen of old Mtb-infected mice at 4 w.p.i. (Fig. 3C and Supplementary Figs. S7 and S8).

**Figure 3.**
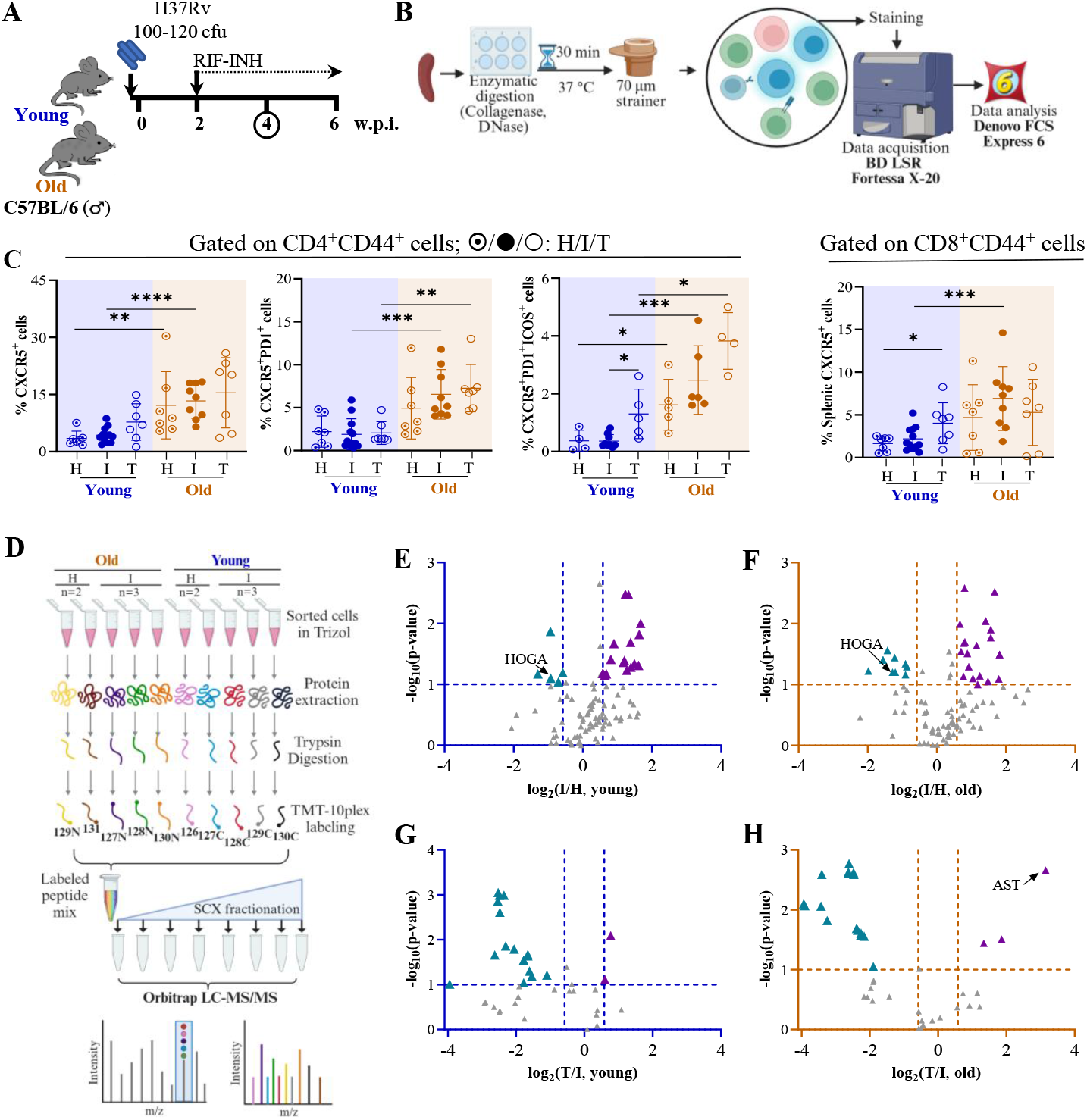
Deregulated splenic CD4^+^CD44^+^ T cell proteome profile observed in old C57BL/6 mice upon Mtb infection and RIF-INH treatment. **A**. Schematic showing Mtb H37Rv infection of male C57BL/6 mice of two age groups (young: 2-4 and old: 17-19 in months) followed by treatment with RIF-INH starting at 2 w.p.i.. **B**. Method adopted for immune cell isolation from the spleen, flow cytometry data acquisition and data analysis. **C**. Frequency of CD4^+^CD44^+^CXCR5^+^, CD4^+^CD44^+^CXCR5^+^PD1^+^ and CD4^+^CD44^+^CXCR5^+^PD1^+^ICOS^+^ and CD8^+^CD44^+^CXCR5^+^ cells in the spleen of C57BL/6 mice at 4 w.p.i./2 w.p.t. are presented. Healthy (H, n=3/age group), Mtb H37Rv infected (I, n= 6-10/age group) and RIF-INH treated (T, n=3/age group). **D**. Schematic representation of method adopted for TMT10plex-T cell proteome analysis (for TMT set-1) by liquid chromatography-mass spectrometry (LC-MS/MS). Volcano plot of splenic CD4^+^CD44^+^ T cell proteome showing significantly deregulated proteins (in -log_10_p-value ≥ 1; log_2_fold change > |0.58|) in (**E**) Mtb infected versus healthy (young), (**F**) Mtb infected versus healthy (old), (**G**) RIF-INH treated versus Mtb infected (young) and (**H**) RIF-INH treated versus Mtb infected (old); 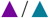: up-/down-regulation; HOGA: 4-hydroxy-2-oxoglutarate aldolase; AST: aspartate aminotransferase. Young mice in blue and old mice in brown; cfu= colony forming unit; w.p.i.= weeks post infection; p-values: * ≤0.05, ** <0.005, *** <0.0005 and **** <0.0001 at 95% confidence interval by Mann-Whitney test. Data is represented as mean ± SD. *Additional details available in Supplementary figures S7, S8, S10 and Supplementary tables S1, S2, S3 and S4*.

In a separate experimental set, at 4 w.p.i., lung Mtb load was found to be similar between young (2M) and aged (31M) male C57BL/6 mice groups (Supplementary Fig. S9). Old and aged mice had overall higher splenic CD4^+^CD44^+^CXCR5^+^PD1^+^, CD4^+^CD44^+^CXCR5^+^PD1^+^ICOS^+^ and CD8^+^CD44^+^CXCR5^+^PD1^+^ cells compared to young mice, irrespective of infection and treatment status (Fig. 3C and Supplementary Fig. S9). The detailed distribution of the immune cells observed in multiple tissues of 2, 4, 17, 19 and 31M old mice groups is tabulated in Supplementary Table S1. The increased abundance of T_FC_-like cells in Mtb-infected old mice suggests dysfunctional T_FH_ cell activity, which may have contributed to compromised pathogen clearance.

### Splenic T cell proteome analysis of old mice upon Mtb infection and anti-TB treatment shows perturbed mitochondrial pathways

To further elucidate the mechanisms underlying the delayed Mtb clearance, we conducted global proteomic profiling of flow-sorted CD4^+^CD44^+^ T cells (Fig. 3D and Supplementary Table S2). In set-I, 109 proteins were identified, and Mtb-infected young and old mice showed 23 and 30 deregulated (p ≤ 0.1, log_2_fold change (log_2_fc): I/H > |0.58|) proteins, respectively (Fig. 3E, 3F and Supplementary Table S3). In set-II, out of 41 identified proteins, ∼20 were deregulated in mice groups receiving treatment (Fig. 3G, 3H and Supplementary Table S4). Proteins related to immune response (e.g., complement, serotransferrin) were suppressed in old mice, potentially reflecting age-related immune decline. Young mice showed few changes, with upregulation of stress response proteins like galectin-1 and superoxide dismutase, possibly reflecting a more robust acute response (Supplementary Table S3). A mitochondrial protein, 4-hydroxy-2-oxoglutarate aldolase (HOGA), was downregulated in the cells of Mtb-infected mice groups irrespective of their age (Supplementary Table S3). Upon Mtb infection, old mice showed an upregulation of Prostaglandin E synthase 3 (log_2_fc: I/H = 1.15), the inhibition of which increases mitochondrial function (Supplementary Table S3). Also, old mice showed elevated levels of mitochondrial AST after treatment (log_2_fc: T/I = 3.15), indicative of greater metabolic adaptation or mitochondrial stress, consistent with increased serum AST and altered hepatic micronutrient status observed earlier (Supplementary Table S4). Old mice showed lower HOGA abundance after treatment, implying that mitochondrial stress persists longer in cells of old mice (Supplementary Table S4). Together, these proteomic insights provide a molecular basis for the observed immune dysregulation, inflammaging, and impaired mycobacterial clearance in older hosts, linking cellular metabolic stress in T cells during Mtb infection and therapy.

## Discussion

As the global demography is shifting towards the older (≥60 years) age groups, it is expected that immunosenescence and lack of an effective TB vaccine for adults will be key drivers for the elevated numbers of geriatric TB patients. Importantly, in countries where TB was contained successfully, the number of new TB cases is increasing, for example, in Kansas City and North Carolina in the United States. The majority of the published reports focused on understanding host-pathogen interactions in Mtb-infected mice of 6-8 weeks (∼2M), representing human age equivalence of 18-20 years^7^. Thus, understanding how the older host handles Mtb infection is crucial for developing appropriate interventions.

TB demonstrates a marked gender disparity, with males showing increased susceptibility (male/female ratio of 1.7 in TB prevalence), which is associated with reduced B cell follicle formation in the lung^12,13,14^. On the contrary, females exhibit increased T cell activity, production of T_H_1-type cytokines, particularly IFN-γ and robust antibody responses^15^. Above 65 years of age, genomic disparities between the sexes become more pronounced, with males having higher innate and pro-inflammatory activity^16^. Thus, we focused on using old male mice to study the outcomes upon Mtb infection and post-treatment.

A low aerosol dose of Mtb infection to young (2-4M), middle-aged (9/12M), old (17-19M) and aged (31M) C57BL/6 mice yielded similar lung mycobacterial burden up to 6 w.p.i.. Whereas earlier reports showed better Mtb control in older mice at earlier time points^8,9,17,18^. For example, Turner et al reported that 18M C57BL/6 mice aerosol infected with Erdman strain had lower lung load at 14 d.p.i., compared to 3M controls^9^. 24M B6D2F1 hybrid mice infected with H37Rv showed lower load at 20 d.p.i. than young 2M controls^8^. Variations in reported tissue Mtb loads in earlier reports and this study may stem from differences in the mouse strain and their sex, Mtb strain and its dose, infection method, and observation timings. Important to highlight that, to the best of our knowledge, this report demonstrates for the first time that old mice showed a compromised Mtb clearance after 2 weeks of RIF-INH treatment compared to younger controls. These results corroborate the reported delayed viral clearance in 17M BALB/c mice infected with PR8 influenza virus^19^.

Aging also impacts hepatic function, affecting metabolism of exogenous molecules, thereby leading to hepatoxicity^20^. Despite their Mtb infection status, old mice showed a higher hepatic Cu/Zn ratio. Accumulation of Cu and Zn in the phagosomes increases bactericidal activity; however, Cu overload is detrimental to the host^21^. Low Zn levels in older mice may impair Cu/Zn superoxide-dismutase function, leading to pro-inflammation and weakened hepatic function^22^. Mtb pathogenesis lowers the available Fe pool and is a typical host response in counteracting Mtb replication in TB patients^23^. Liver Fe level was reduced in young mice post treatment; however, higher hepatic Fe pool in the treated old mice may help Mtb for its growth and multiplication. Impaired liver function, indicated by higher AST levels, can lead to reduced drug metabolism and elimination. Elevated serum AST and ALT support the concept that aging is linked to increased liver stress, which may contribute to altered responses to liver insults such as drug treatment post-infection.

Immune cells like CD4^+^ T cells play a critical role in controlling Mtb, and their function might be partially impaired in older hosts^24^. Higher lung mycobacterial load was predicted in 18M mice at 6 w.p.i. compared to 3M mice, due to a delayed CD4^+^ T cell response^24,25^. We observed that Mtb-infected old C57BL/6 mice had similar T_FH_-like cells in the lung and a higher frequency in the spleen at 4 w.p.i. compared to younger controls. However, a higher proportion of T_FC_-like cells in the lung and spleen was observed in Mtb-infected old mice. The contribution of T_H_1 cells in TB is well established. However, our understanding of T follicular cells, which link humoral and adaptive immunity, during Mtb infection is limited^26^. T_FC_ cells suppress T_FH_ cell helper function and antibody responses through multiple mechanisms, inhibiting antibody production to maintain self-tolerance and regulate immune responses^27,28^. Age is reported to influence T_FH_ cell function and differentiation negatively, compromising the host responses^29^. Localization of T_FH_-like cells has been demonstrated within protective granuloma-associated lymphoid tissue in lungs of 2M C57BL/6 mice which mediates Mtb control^30^.

Understanding tissue level T cell distribution in Mtb-infected old mice has been attempted earlier; however, limited reports captured the changes during TB treatment^8,9,17,24,26^. At 2 w.p.t., young mice showed increased splenic T_FH_-like cells, which remained similar in old mice, suggesting the inability of the older host to elicit a proper T_FH_ response. Compromised Mtb clearance in old mice might be due to the impaired T_FH_ cell functionality, which was reported earlier in >20M old C57BL/6 mice^29,31^.

The splenic cell proteome showed significant variation in the abundance of key mitochondrial enzymes, e.g., HOGA and AAT, involved in the catabolism of hydroxyproline (Hyp), a product of protein degradation (Fig. 4)^32^. AAT catalyzes 4-hydroxy glutamate (4-OH-Glu) to 4-hydroxy-2-oxo-glutarate (HOG), which is converted into pyruvate and glyoxylate using HOGA^32^. A higher abundance of AAT and lower HOGA levels in the splenic cells of Mtb-infected old mice might lead to HOG accumulation. HOG inhibits glyoxylate reductase, which contributes to excessive oxalate formation independent of the lactate dehydrogenase pathway and is reported to cause mitochondrial dysfunction^33,34^. We found higher levels of Prostaglandin E synthase-3 in Mtb-infected old mice, which is reported to negatively affect the proliferation of alveolar macrophages (AMs)^35^. Also, inhibition of prostaglandin E2 receptor enhanced mitochondrial function in AMs of 18-22M mice^35^. Reports demonstrated the potential of mitochondrial-based interventions to enhance aged CD4^+^ T cell functionality with implications for improving immune responses in geriatric TB patients^36,37^. This study has a few limitations. Due to a limited number of lung T_FH_ cells, we could not profile their proteome level changes and focused on the splenic CD4^+^CD44^+^ T cells. Mouse models in TB research have limitations for extrapolating findings to humans due to differences in disease pathology, lacking key features like granuloma liquefaction, cavitation, and fibrosis. Also, mice exhibit different metabolic and aging patterns compared to humans, with distinct changes in organ metabolism, immune responses, and biomarkers of aging.

**Figure 4.**
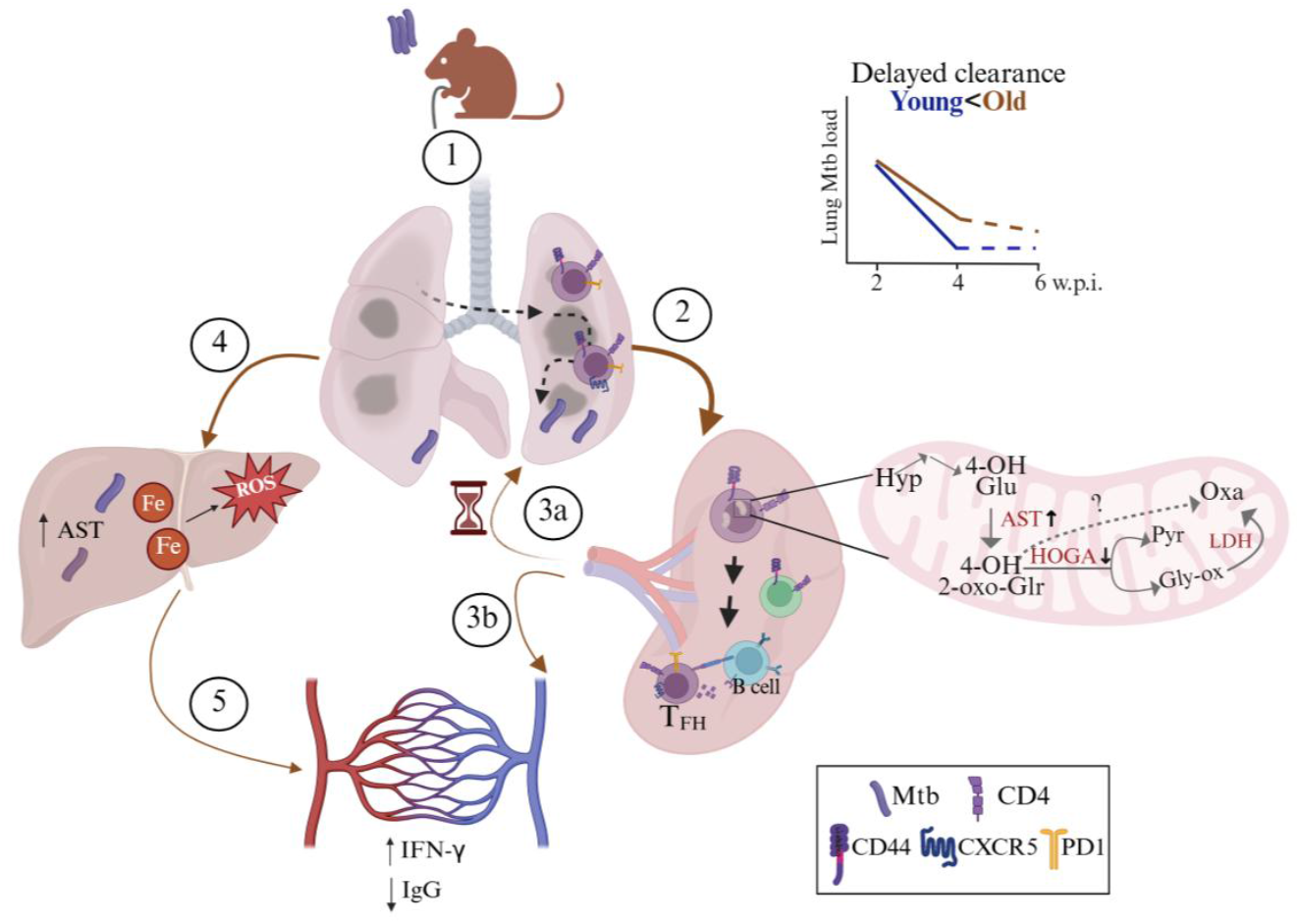
Old C57BL/6 mice showed a compromised lung *Mycobacterium tuberculosis* (Mtb) H37Rv clearance in the early time points of rifampicin and isoniazid treatment. Illustrative summary of the results observed in the study. (**1**) Following Mtb infection via aerosol challenge, (**2**) antigen presentation takes place in the spleen. (**3a**) T cells take longer to reach the inflamed lung of old mice. (**3b**) Increased levels of proinflammatory cytokines: interferon-gamma (IFN-γ) and decreased Mtb-specific IgG levels observed in Mtb-infected old mice. (**4**) Mtb disseminates to the liver via hepatic artery where iron (Fe) was observed to be accumulated along with higher Aspartate aminotransferase (AST) activity, suggestive of higher reactive oxygen species (ROS) generation (**5**) which also in turn could increase circulatory proinflammatory mediators. Old mice had low splenic T follicular helper (T<SUB>FH </SUB>: CD4^+^CD44^+^CXCR5^+^PD1^+^) cells, and presented a delayed lung Mtb clearance at 2 weeks post treatment. Also, proteome analysis of splenic CD4^+^CD44^+^ T cells of old mice showed alterations in mitochondrial proteins (HOGA: 4-hydroxy-2-oxoglutarate aldolase and AST) which are related to hydroxyproline (Hyp) degradation pathway. Increased AST and decreased HOGA might lead to buildup of 4-hydroxy-2-oxo-glutarate (4-OH-2-oxo-Glr) from glutamate (Glu), directing the reaction to oxalate (Oxa) via pathway independent of lactate dehydrogenase (LDH), pyruvate (Pyr) and glyoxylate (Gly-ox). Figure created with BioRender.com

In summary, this study revealed an impaired lung Mtb clearance in old C57BL/6 mice receiving RIF-INH, mainly due to inflammaging, delayed T_H_1 responses, higher T_FC_ cell frequency and mitochondrial dysfunction in T cells. The findings from this study help us to better understand the effect of immunosenescence on poorer treatment outcomes in geriatric TB patients.

## Materials and Methods

### Experimental groups, Mtb infection and treatment

Procedures adopted in this study were performed following the recommended Guidelines for Care and Use of Laboratory Animals and approved by the Animal Ethics Committee of ICGEB-New Delhi (Ref No. ICGEB/IAEC/07032020/TH-10). C57BL/6 mice of different age groups (2,4,9,12,17,19 and 31 months) were procured from the ICGEB bio-experimentation facility. These mice were transferred to the BSL-III Tuberculosis Aerosol Challenge Facility (TACF) for Mtb infection experiments. Mice were fed a chow diet, water ad libitum and maintained at 20-22ºC; 45-60% humidity with a 12h light/dark cycle. Mice were weight-adjusted, randomly grouped and after 1 week of acclimatization in TACF, infected with a low dose (100-120 cfu) of Mtb H37Rv strain in the Madison aerosol exposure chamber. After 2 w.p.i., a subset of mice received RIF and INH (100 mg/L each) in drinking water and fresh drug solutions were replaced every 48 hours.

### Cfu assay

Tissues (lung, spleen and liver) were homogenized in PBS (1 mL) and plated on 7H11 agar supplemented with Middlebrook 10% OADC. These plates were incubated at 37ºC and bacterial colonies were enumerated 3 weeks after.

### Histopathology

Formalin-fixed lungs were paraffin-embedded, sectioned and stained with hematoxylin and eosin, followed by capturing images using Nikon Eclipse TS2. The slides were scored based on the severity of inflammation, granuloma formation and necrosis.

### Liver micronutrient profiling

Liver (∼50 mg) was subjected to microwave digestion post addition of H_2_O_2_ and HNO_3_ (1:4). These were diluted and introduced to iCAP™-TQ ICP-MS using argon as carrier. Data acquisition was done using Qtegra (Thermo Scientific, version: 2.10.4345.39) along with running different concentrations of multielement standard mix (Supelco, #92091).

### ALT and AST assay

Serum ALT and AST activities were measured using kits (Cayman Chemical, #700260 and #701640) and steps provided by the manufacturer.

### Immune cell isolation and flow cytometry

Harvested tissues were incubated with collagenase D (1 µg/mL) and DNase I (0.5 mg/mL) and stained with: CD3-BV510, CD4-BV785, CD8-AF700, CD44-BV605, CXCR5-PE/Dazzle, PD1-PE-Cy7, KLRG1-BV421, CD153-PE, CD25-AF488 and CD127-BV650 and live dead-APC-Cy7 on ice for 45 min. After washing with buffer (0.2% FBS in 1× PBS), cells were treated with fixation and permeabilization buffer and kept on ice for 30 min. Intracellular staining was done with FoxP3-AF647 and Tbet-BV711 and incubated at RT for 45 min, followed by washing. Data (∼1 million events/sample) was acquired using BD LSRFortessa X-20 and analyzed using FCS Express (DeNovo software, version 6).

Another set of immune cells were stained with: CD3-FITC, CD4-SB600, CD8a-BV421, CD44-BV510, CXCR5-PE, ICOS-PerCP/Cy5.5, PD1-PE-Cy7 and live dead-APC-Cy7 on ice for 45 min. After washing with buffer (3% FBS in 1× PBS), splenic CD4^+^CD44^+^ T cells were sorted using BD FACS Aria III Fusion. Data was analyzed using FlowJo (BD Biosciences, version 10.8.1).

### Serum cytokine estimation

Two times diluted serum samples were used for estimating cytokines using capture bead-based assay LEGENDplex (BioLegend) along with the standard (BioLegend, #740371, lot no. B330381). The samples were acquired on BD LSRFortressa X-20 using PE and APC as the reporter and bead classification channel, respectively. Data was analyzed using Qognit software.

### Serum IgG antibody quantification

ELISA plate was coated with Mtb H37Rv whole cell lysate (10 µg/mL) and incubated overnight at 4°C. Then, wash buffer (0.05% Tween-20 in 1× PBS) was added, followed by blocking in 3% BSA in 1× PBS containing 0.05% Tween-20 and incubated at 37°C for 2h at 180 rpm. After blotting the plate dry, sera were serially diluted four-fold in 0.1% BSA in 1× PBS containing 0.05% Tween-20, and 100 µL was added per well and incubated overnight at 4°C. The plate was washed thrice followed by incubation with anti-mouse IgG (Southern Biotech, #1031-05) at 37°C for 2h at 180 rpm. The plate was washed six times and developed by adding o-phenylenediamine dihydrochloride, incubated at RT for 5 min. The reaction was terminated by adding 1N HCl and the absorbance was measured at 490 nm.

### Splenic CD4^+^CD44^+^ cell proteome analysis

Proteins were isolated using TRIzol reagent from sorted splenic CD4^+^CD44^+^ cells and quantified using BCA Protein Assay. An equal amount (∼10 µg) of proteins from a set of samples (n=10) was taken for TMT10plex (Thermo Scientific, #90113). TMT-labelled peptides from all samples were pooled, dried and subjected to strong cation exchange chromatography and eluted using ammonium formate. Later, these fractions (n = 5) were cleaned up using C18 spin columns and dried, resuspended in formic acid (0.1%, 20 μL) and injected into an Orbitrap Fusion Lumos Tribrid Mass Spectrometer connected to a nano-LC 1200. Data was analysed with Proteome Discoverer version 2.3 using *Mus musculus* (ID: UP000000589; 55,087 protein count). Proteins qualifying the parameters [log_2_ fold change ≥ ± 0.58; log_10_p-value ≥1; false discovery rate<0.05] were selected as important deregulated molecules.

### Statistical analysis

Data are presented as mean ± standard deviation. Details regarding the sample size and statistical analysis have been provided in the figure legends.

## Acknowledgements

FP was supported with a Junior and Senior Research fellowship from the Department of Biotechnology (DBT), Government of India. SC was a Shyama Prasad Mukherjee Fellow supported by the Council of Scientific and Industrial Research, Government of India. A Junior Research Fellowship from the DBT supports AG. The core support from the International Centre for Genetic Engineering and Biotechnology (ICGEB), New Delhi, to RKN is highly acknowledged. Haripriya Priyadarsini, Adyasha Sarangi, Nidhi Yadav and Mothe Sravya are acknowledged for their help and support during the mouse experiments. Help from the staff of the bio-experimentation facility, BSL-III Tuberculosis Aerosol Challenge Facility (TACF) at ICGEB-New Delhi, Shilpa Sharma and Jaya Baranwal, is acknowledged. DBT supports the TACF. We thank Dr. Anmol Chandele for her insightful comments and suggestions for revising the manuscript. Part of these research findings were presented in Keystone Symposia on Molecular Basis of Healthy Aging in 2023; World Conference on Lung Health 2022, organized by The Union, and the 2^nd^ biennial meeting on Sex Differences in the Immune Systems Conference in 2022, organized by Trinity College Dublin & Whitehead Institute. Some figures have been created with BioRender. The authors declare no competing financial interests.

## Author Contributions

FP and RKN conceptualized the project. FP and SC carried out the mouse experiments and cytokine profiling. FP performed the flow cytometry data acquisition and liver micronutrient and proteomics sample preparation. AG carried out antibody titer, ELISA experiments and helped in proteomic data analyses. ShiC calculated histopathological scores. FP and RKN analyzed the data, wrote the original draft and revised it with the suggestions received from all the co-authors. RKN acquired funding, shared resources and administered the project. All co-authors reviewed and approved the manuscript.

## Declaration of interests

The authors declare no competing interests.

## Supporting Information

The article contains supplemental information.

## Notes

### Competing Interest Statement

The authors have declared no competing interest.

### Summary of Updates

The paper has been revised with new experimental sets. One author has been added.

