## Supplementary Figures S1 to S10; Supplementary Tables S1 to S4. for "Host immunosenescence compromises *Mycobacterium tuberculosis* clearance"

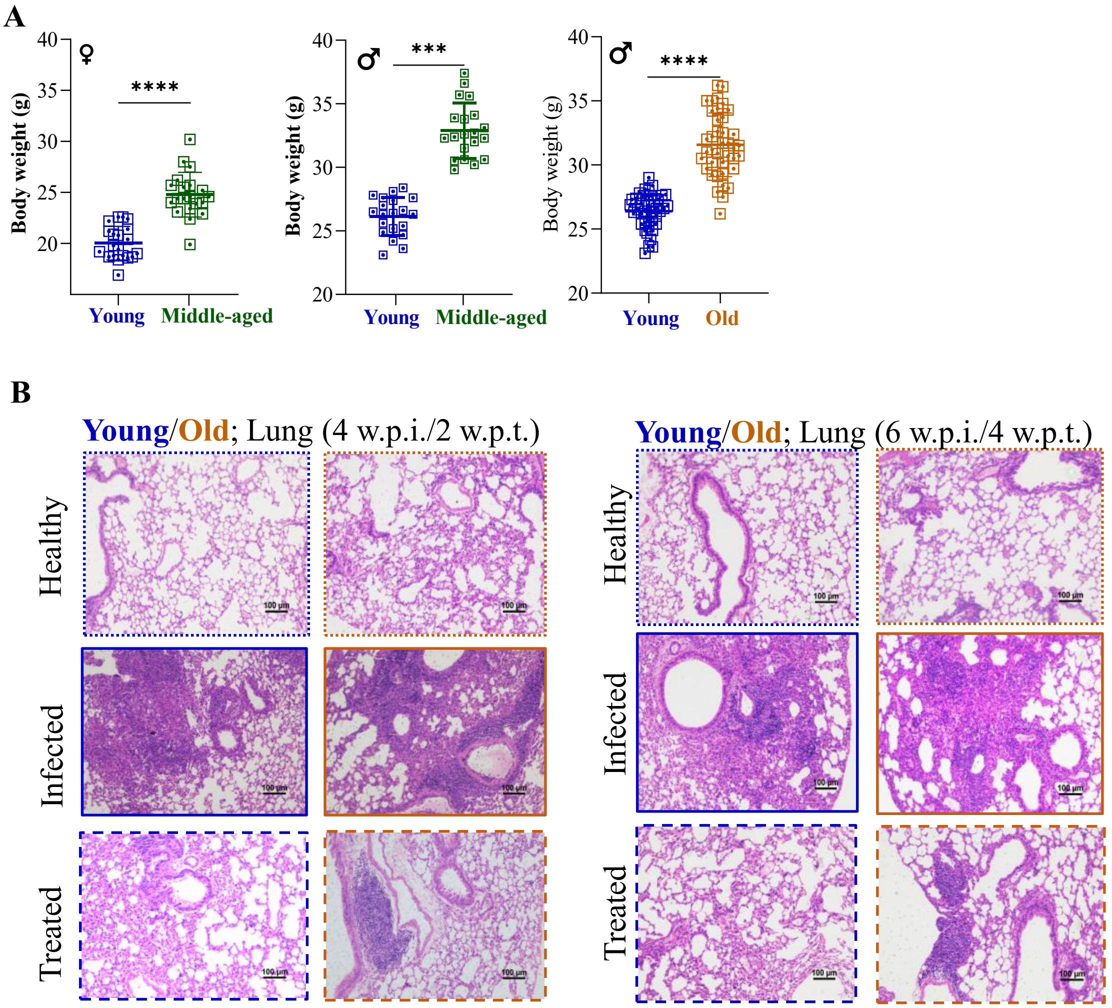


**Supplementary Fig. S1: Old C57BL/6 mice receiving two weeks of RIF-INH treatment showed delayed lung Mtb clearance**. **A**. Body weight (in gram) of mice (n=40/age group) prior to infection (day -5). **B**. Lung histopathological images (hematoxylin and eosin; H&E stained, 100× magnification, scale bar=100 μm) at 4 w.p.i./2 w.p.t. and 6 w.p.i./4 w.p.t. Young (2-4 months) in blue, middle-aged (9-12 months) in green and old (17-19 months) mice in brown; w.p.i.= weeks post infection; w.p.t.= weeks post treatment; p-values: *** <0.0005 and **** <0.0001 at 95% confidence interval by Mann-Whitney test. Data is represented as mean ± SD. *This figure is an extension of Figs. 1 and 2.*

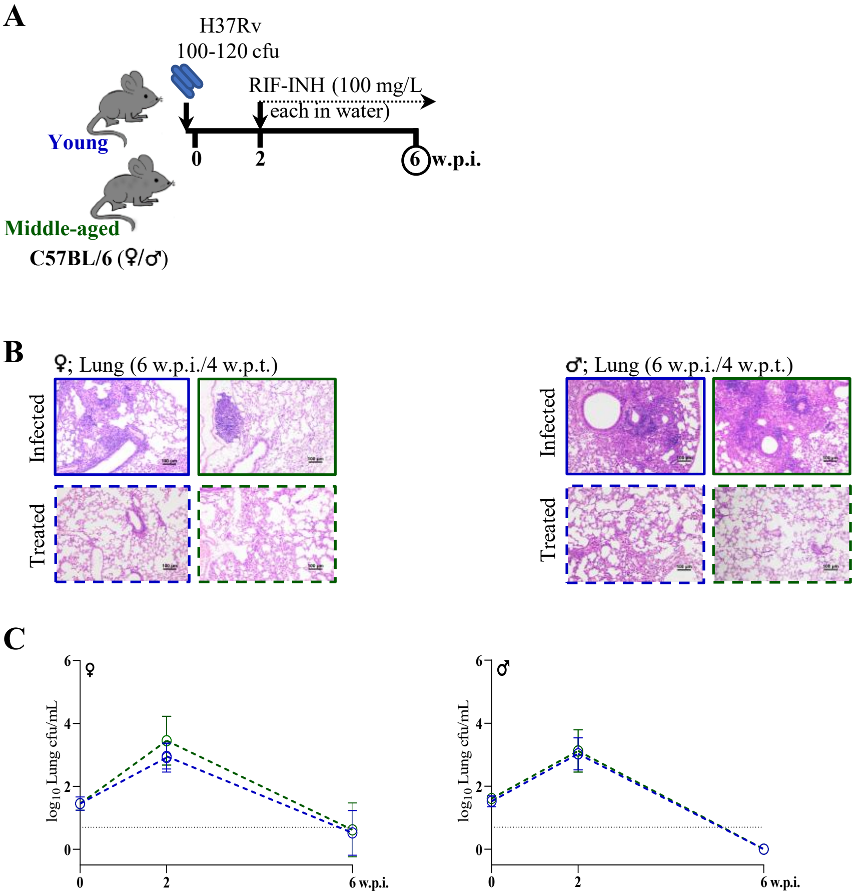


**Supplementary Fig. S2: Middle-aged C57BL/6 mice showed similar treatment outcomes post Mtb infection compared to younger mice irrespective of gender**. **A**. Schematic of the experimental design for Mtb H37Rv infection and RIF-INH treatment of C57BL/6 mice (female: age groups of 2 and 9 months and male: age groups of 2 and 12 months; M). **B**. Lung histopathological images (hematoxylin and eosin: H&E stained, 100× magnification, scale bar=100 µm) of infected (I) and RIF-INH treated (T) groups at 6 w.p.i./4 w.p.t. **C**. Lung bacterial burden (in $log_{10}$cfu/mL) in mice belonging to treated group (at 4 w.p.t.; treatment started at 2 w.p.i.) groups; n=5/timepoint/age group/condition; dashed horizontal line represents limit of detection. Young mice in blue and middle-aged mice in green; cfu: colony forming unit; w.p.i.: weeks post infection; w.p.t.: weeks post treatment. Data is represented as mean ± SD. *This figure is an extension of Fig. 1.*

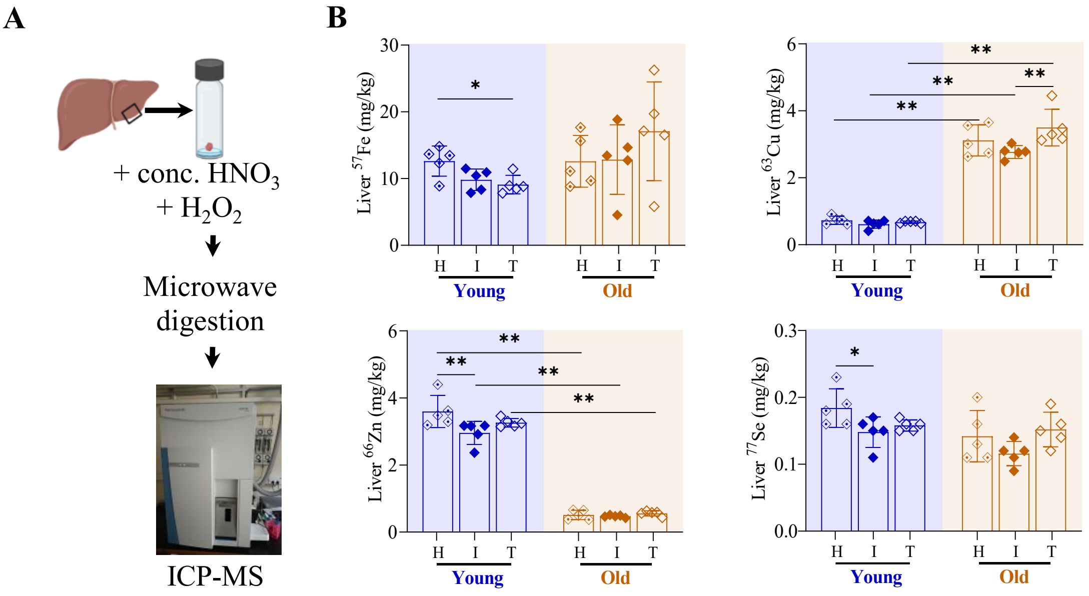


**Supplementary Fig. S3: Impaired Mtb clearance in old C57BL/6 mice potentially correlates with abnormal levels of liver micronutrients**. **A**. Method adopted for liver micronutrient profiling by inductive coupled plasma mass spectrometry (ICP-MS). **B**. Abundance of mice liver micronutrients (in milligram per kilogram): iron- $^{57}$Fe, copper- $^{63}$Cu, zinc- $^{66}$Zn, and selenium- $^{77}$Se of H, I and T groups at 4 w.p.i./2 w.p.t.; n=5/age group/condition. p-values: * $\leq$0.05 and ** <0.005 at 95% confidence interval by Mann-Whitney test. Data is represented as mean ± SD. Young (2 months) in blue and old (17 months) mice in brown.

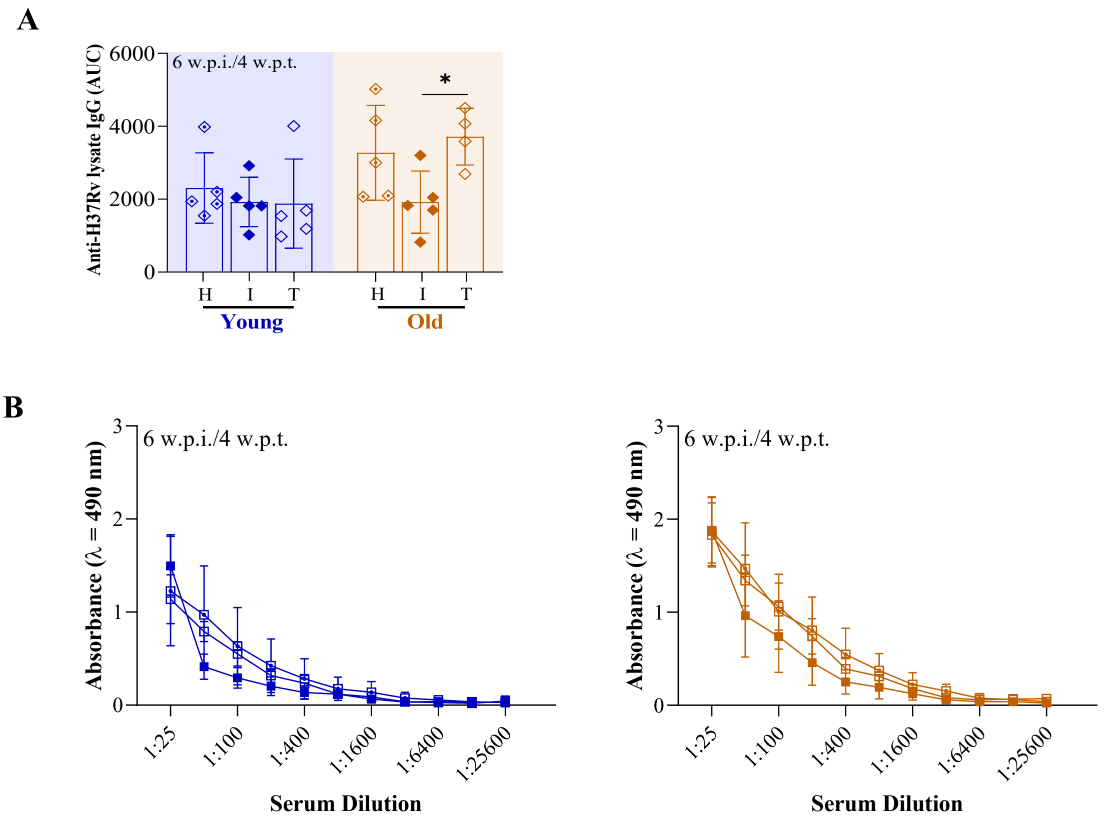


**Supplementary Fig. S4: Serum anti-Mtb IgG was higher in old mice after RIF-INH treatment**. **A**. Anti-H37Rv lysate IgG estimation with area under the curve (AUC) in the circulation of healthy (H), Mtb H37Rv infected (I) and RIF-INH treated (T) C57BL/6 mice at 6 w.p.i./4 w.p.t.. **B**. End-point serum dilutions to monitor anti-IgG titers in mice groups at 6 w.p.i./4 w.p.t.; n = 5/time point/age group/condition (only old-T had n=4 at 4 w.p.t.). Young (2 months) in blue and old (17 months) mice in brown; p-value: * ≤0.05 at 95% confidence interval by Mann Whitney test. Data is represented as mean ± SD. *This figure is an extension of Fig. 2.*

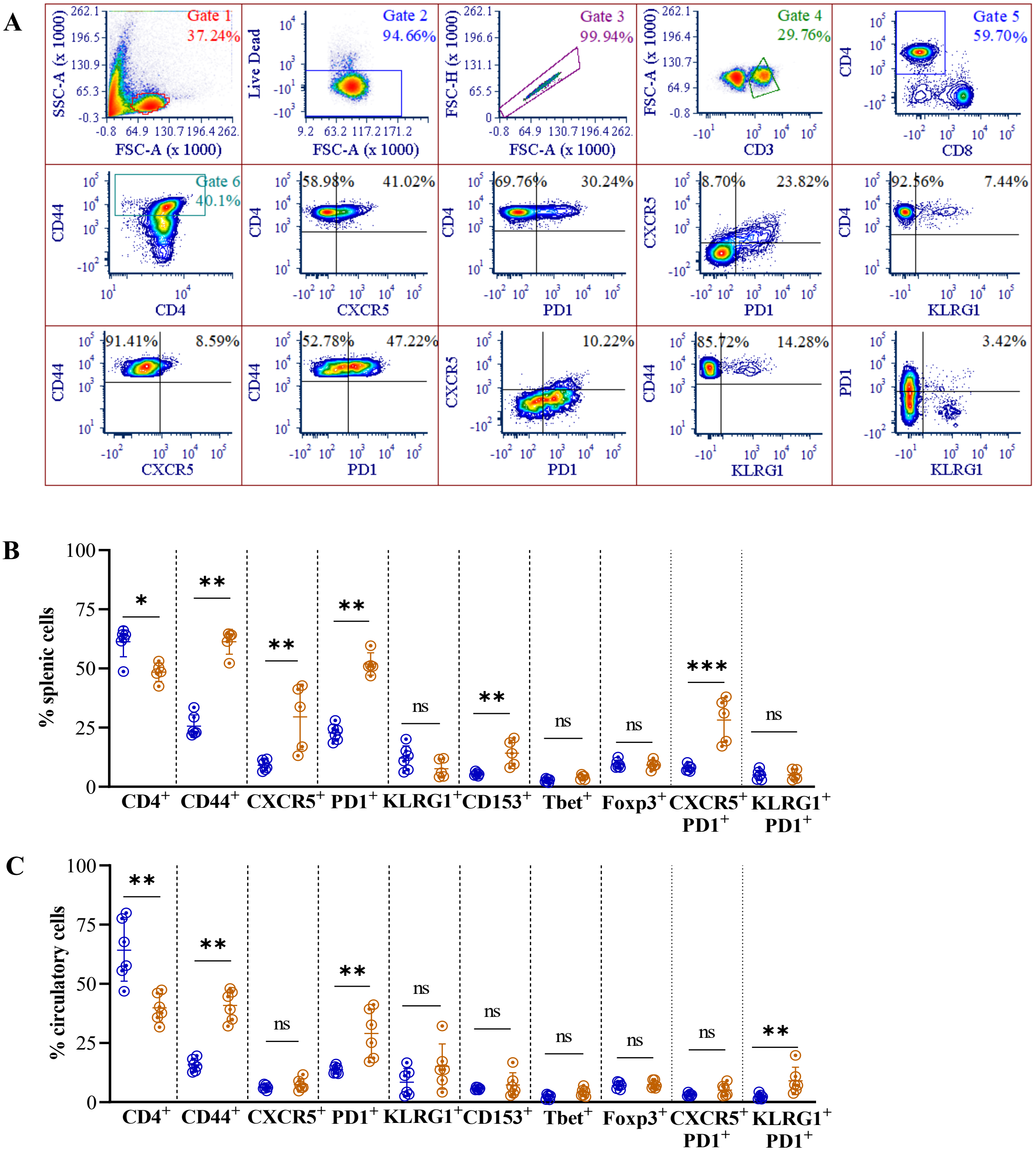

**Supplementary Fig. S5: Splenic and circulatory $CD4^{+}$ T cell subsets frequency altered in old C57BL/6 mice**. **A**. Gating strategy in the lung of a representative mouse adopted to monitor the frequency of multiple $CD4^{+}$ T cell subsets; *data analyzed using FCS Express (version 6)*. Frequency of $CD4^{+}$, $CD4^{+}CD44^{+}$, $CD4^{+}CXCR5^{+}$, $CD4^{+}PD1^{+}$, $CD4^{+}KLRG1^{+}$, $CD4^{+}CD153^{+}$, $CD4^{+}Tbet^{+}$, $CD4^{+}CD25^{+}CD127^{-}FoxP3^{+}$, $CD4^{+}CXCR5^{+}PD1^{+}$ and $CD4^{+}KLRG1^{+}PD1^{+}$ cells in the (**B**) spleen and (**C**) circulation of healthy mice are presented. n = 6 for young, n = 5-6 for old mice; Young (2 months) in blue and old (17 months) mice in brown; p-values: ns= non-significant, * ≤0.05, ** <0.005 and *** <0.0005 at 95% confidence interval by Mann-Whitney test. Data is represented as mean ± SD.

A

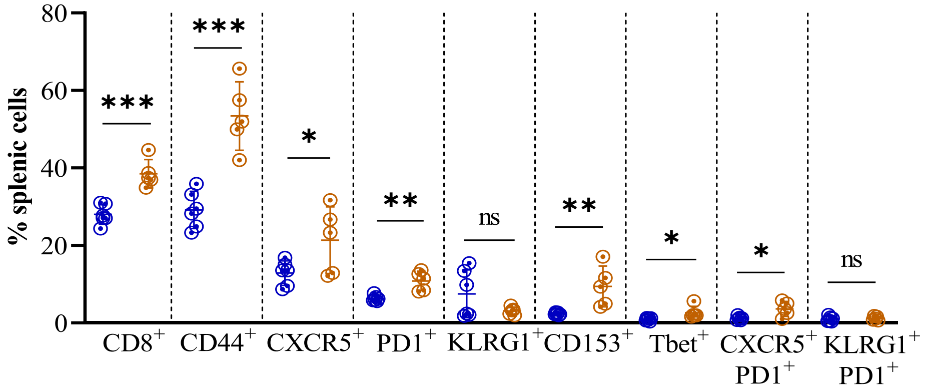


B

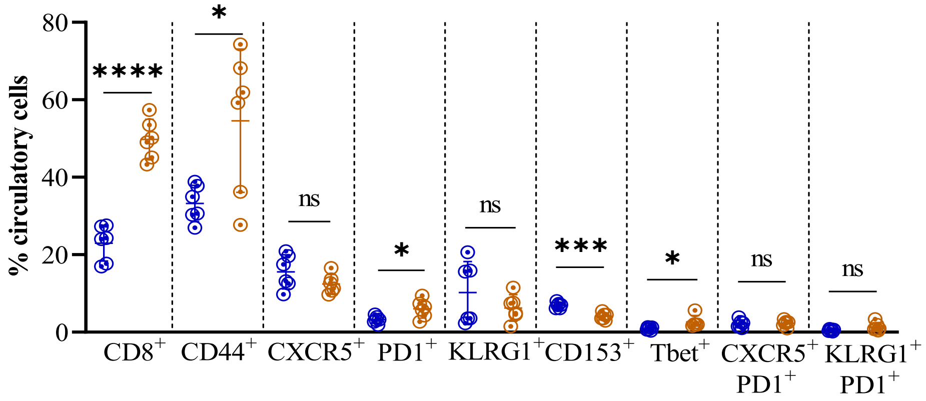


**Supplementary Fig. S6: Splenic and circulatory CD8$^{+}$ cells subsets altered in old mice**. Frequency of CD8$^{+}$, CD8$^{+}$CD44$^{+}$, CD8$^{+}$CXCR5$^{+}$, CD8$^{+}$PD1$^{+}$, CD8$^{+}$KLRG1$^{+}$, CD8$^{+}$CD153$^{+}$, CD8$^{+}$Tbet$^{+}$, CD8$^{+}$CXCR5$^{+}$PD1$^{+}$, CD8$^{+}$KLRG1$^{+}$PD1$^{+}$cells in the (**A**) spleen and (**B**) circulation of healthy young (n=6) and old (n=5-6) mice are presented. Young (2 months) in blue and old (17 months) mice in brown; p-values: ns = non-significant, * ≤0.05, ** <0.005, *** <0.0005 and **** <0.0001 at 95% confidence interval by Mann Whitney test. Data is represented as mean ± SD.

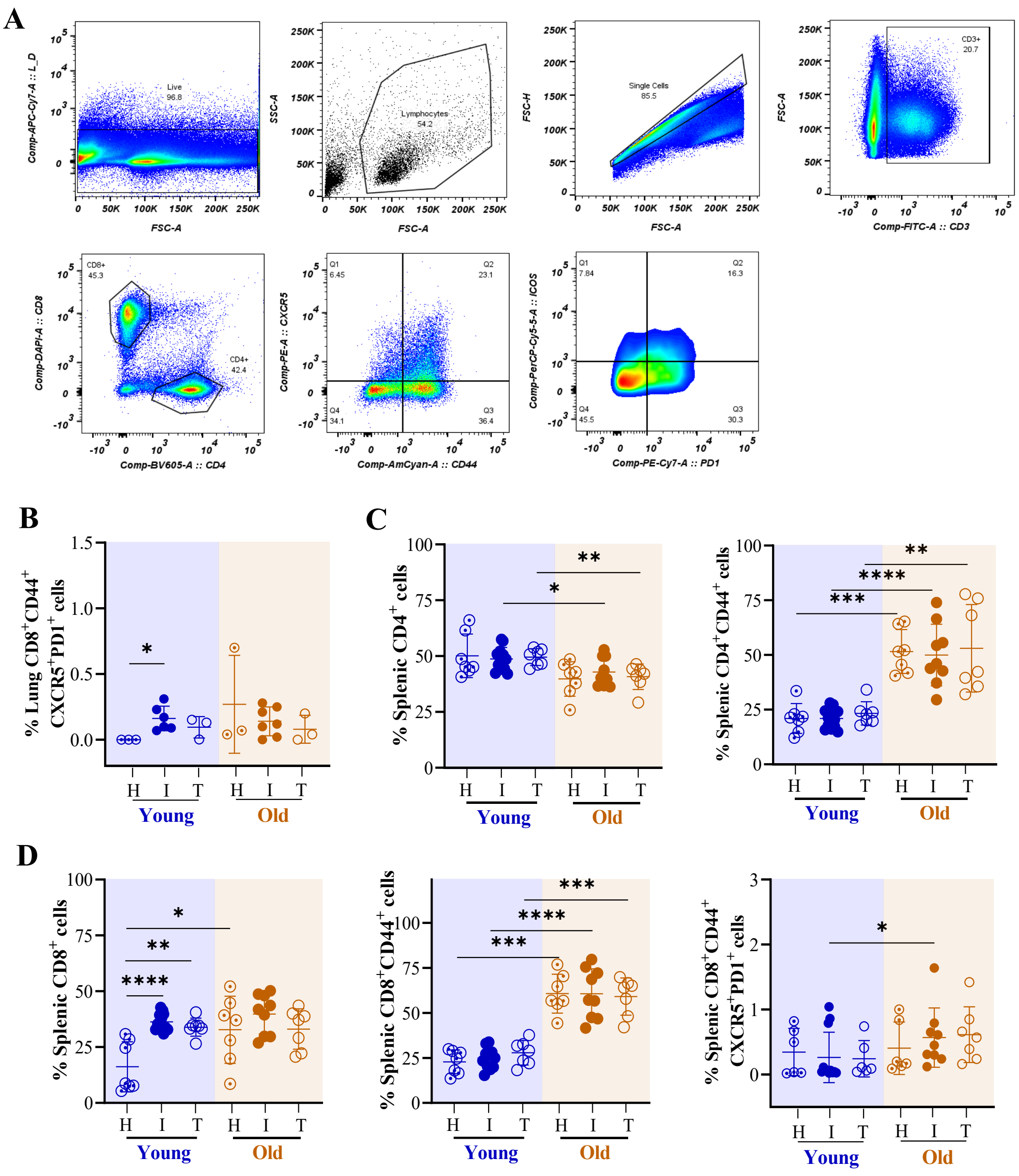


**Supplementary Fig. S7: Higher lung and splenic CD8$^+$CD44$^+$CXCR5$^+$ cells observed in Mtb infected old mice**. **A**. Gating strategy of a representative mouse adopted to monitor the frequency of T cell subsets in the lung of C57BL/6 mice; *data analyzed using FlowJo (version 10.8).* **B**. Frequency of lung CD8$^+$CD44$^+$CXCR5$^+$PD1$^+$ cells of healthy (H, n=3/age group), Mtb H37Rv infected (I, n= 9-10/age group) and RIF-INH treated (T, n=3/age group) C57BL/6 mice at 4 w.p.i./2 w.p.t.. Frequency of splenic (**C**) CD4$^+$ and CD4$^+$CD44$^+$ (**D**) CD8$^+$, CD8$^+$CD44$^+$ and CD8$^+$CD44$^+$CXCR5$^+$PD1$^+$ cells of healthy (H, n=6-7/age group), Mtb H37Rv infected (I, n=9-12/age group) and RIF-INH treated (T, n=6-7/age group) C57BL/6 mice at 4 w.p.i./2 w.p.t. Young (2-4 months) in blue and old (17-19 months) mice in brown; p-values: * ≤0.05, ** <0.005, *** <0.0005 and **** <0.0001 at 95% confidence interval by Mann Whitney test. Data is represented as mean ± SD. *This figure is an extension of Fig. 3.*

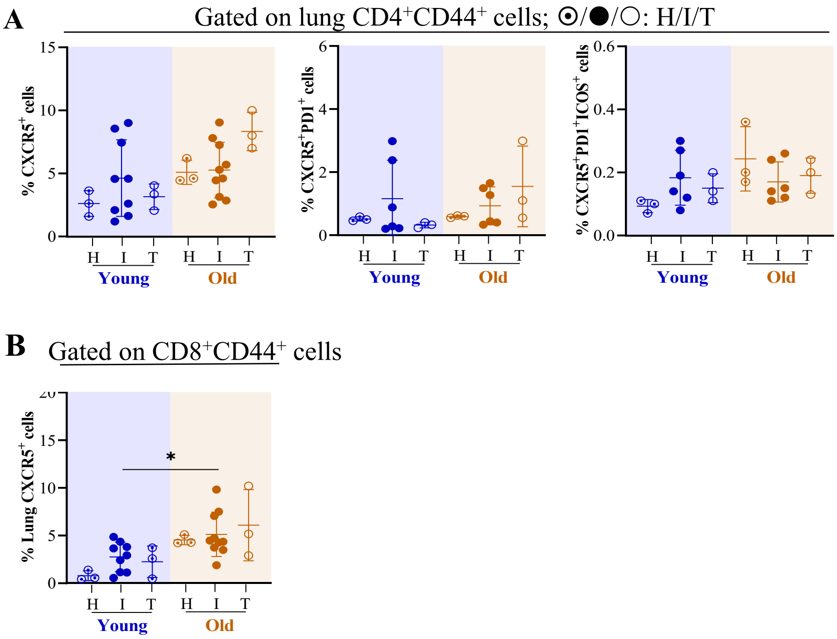


**Supplementary Fig. S8: Lung CD8$^+$CD44$^+$CXCR5$^+$ cells were higher in old C57BL/6 mice post Mtb infection**. **A**. Frequency of CD4$^+$CD44$^+$CXCR5$^+$, CD4$^+$CD44$^+$CXCR5$^+$PD1$^+$ and CD4$^+$CD44$^+$CXCR5$^+$PD1$^+$ICOS$^+$ cells in the lungs of C57BL/6 mice at 4 w.p.i./2 w.p.t. are presented. **B**. Frequency of CD8$^+$CD44$^+$CXCR5$^+$ cells in the lungs and spleen at 4 w.p.i./2 w.p.t. are presented. Healthy (H, n=3/age group), Mtb H37Rv infected (I, n= 6-10/age group) and RIF-INH treated (T, n=3/age group). Young (2-4 months) in blue and old (17-19 months) mice in brown; p-value: * ≤0.05 at 95% confidence interval by Mann-Whitney test. Data is represented as mean ± SD. *This figure is an extension of Fig. 3.*

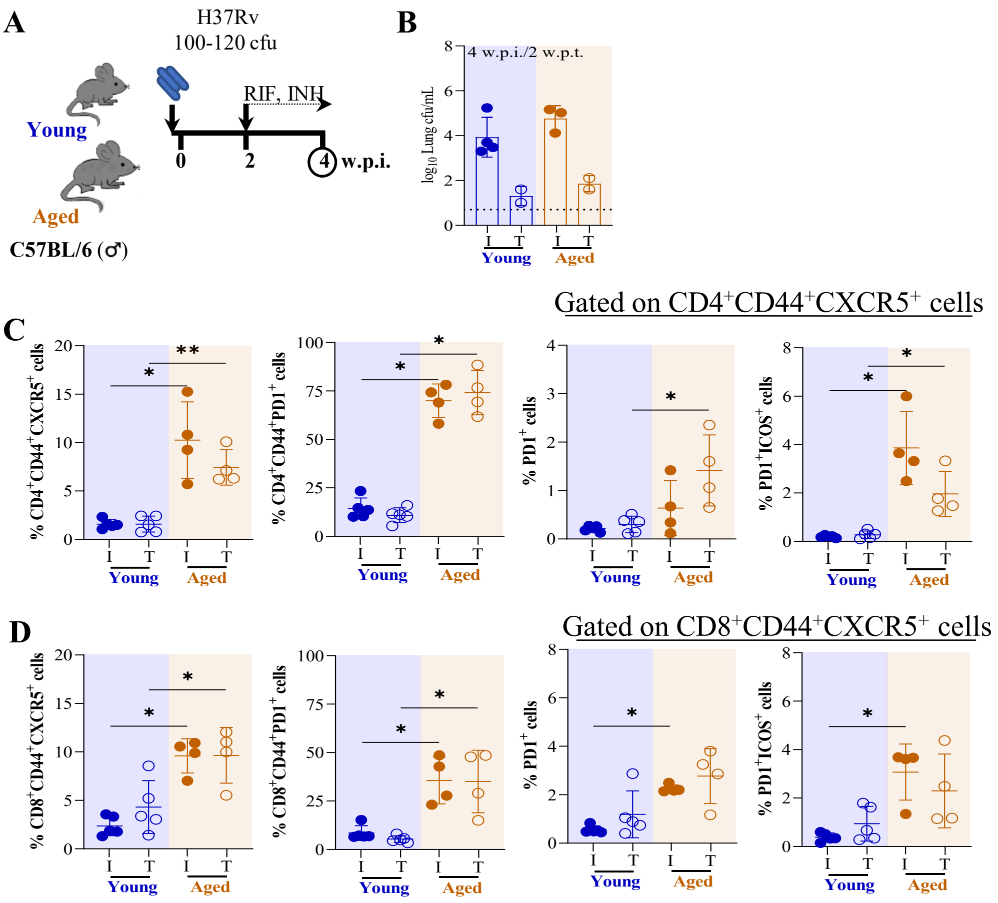


**Supplementary Fig. S9: Higher splenic CD4$^+$CD44$^+$CXCR5$^+$ cells observed in aged mice**. **A**. Schematic of the experimental design for Mtb H37Rv infection and RIF-INH treatment of male C57BL/6 mice. **B**. Lung bacterial burden (in log$_{10}$cfu/mL) of Mtb infected mice (at 4 w.p.i.) and RIF-INH treated mice (at 2 w.p.t.; treatment started at 2 w.p.i.); dashed horizontal line represents limit of detection (LOD); Infected (I) mice: n=3-4 for young and old; Treated (T) mice: n=2 for young and old; rest cfu data points could not be collected due to plate contamination. **C**. Frequency of splenic CD4$^+$CD44$^+$CXCR5$^+$, CD4$^+$CD44$^+$PD1$^+$, CD4$^+$CD44$^+$CXCR5$^+$PD1$^+$ and CD4$^+$CD44$^+$CXCR5$^+$PD1$^+$ICOS$^+$ cells of I and T groups of mice at 4 w.p.i./2 w.p.t. **D**. Frequency of splenic CD8$^+$CD44$^+$CXCR5$^+$, CD8$^+$CD44$^+$PD1$^+$, CD8$^+$CD44$^+$CXCR5$^+$PD1$^+$ and CD8$^+$CD44$^+$CXCR5$^+$PD1$^+$ICOS$^+$ cells of I and T groups of mice at 4 w.p.i./2 w.p.t. n=4-5/condition; Young (2 months) in blue and aged (31 months) mice in brown; p-values: * ≤0.05 and ** <0.005 at 95% confidence interval by Mann Whitney test. Data is represented as mean ± SD.

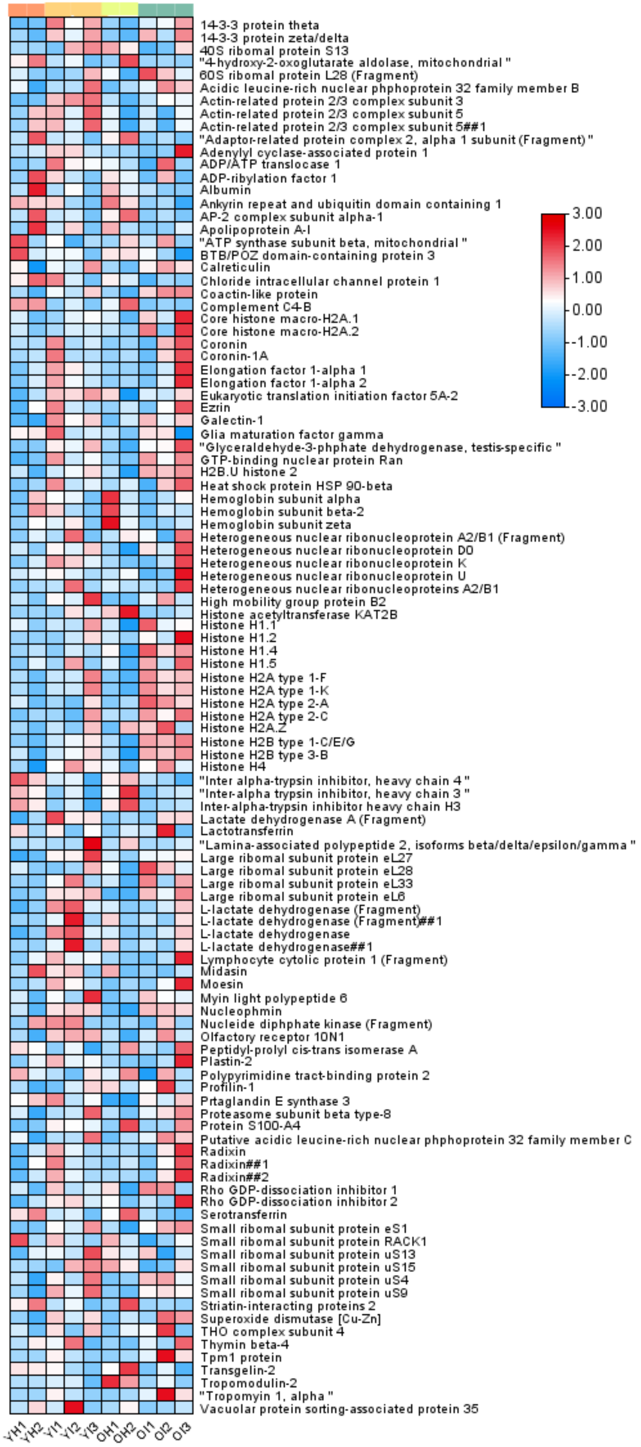


**Supplementary Fig. S10: Proteome distribution of CD4$^+$CD44$^+$ T cells changed upon Mtb infection**. Heatmap showing abundance of identified splenic CD4$^+$CD44$^+$ T cell proteins obtained from TMT10plex experiment using healthy (H) and Mtb H37Rv infected (I) C57BL/6 mice groups (Y: young, 4 months) and (O: old, 19 months). Each row presents abundance of individual protein. 1,2,3 shows the biological replicates. Colour intensity indicates the abundance values. *This figure is an extension of Fig. 3.*

**Supplementary Table S1: Compiled results of the study**. Distribution of different immune cells in young (2M, 4M), old (17M, 19M) and aged (31M) male C57BL/6 mice in (**A**) spleen of healthy (H), Mtb H37Rv infected (I) and RIF-INH treated (T) groups at 4 w.p.i./2 w.p.t. and in (**B**) lung of Mtb infected mice at 4 w.p.i.; M: months. *This table is an extension of Fig. 3.*

**A**

| Splenic cells | Condition | 2M | 4M | 17M | 19M | 31M |
|---|---|---|---|---|---|---|
| $CD4^+$ | H | 58±9 | 43±2 | 45±4 | 37±8 | N/A |
| | I | 55±2 | 46±3 | 47±6 | 40±5 | 42±5 |
| | T | 49±5 | 49±4 | 43±4 | 41±7 | 43±9 |
| $CD4^+CD44^+$ | H | 25±4 | 17±4 | 61±5 | 49±10 | N/A |
| | I | 23±2 | 20±5 | 65±9 | 42±8 | 88±5 |
| | T | 21±2 | 23±6 | 73±5 | 37±4 | 86±6 |
| $CD4^+CD44^+CXCR5^+$ | H | 3±0.5 | 4±2.6 | 6±1.4 | 14±10 | N/A |
| | I | 4±0.3 | 4±2.3 | 8±1.5 | 16±3 | 10±4 |
| | T | 2±1.6 | 10±3 | 6±3.2 | 23±4 | 7.4±1.8 |
| $CD4^+CD44^+CXCR5^+PD1^+$ | H | 3.8±1.3 | 1±0.7 | 8±4.7 | 3.4±2 | N/A |
| | I | 4.5±1.4 | 1±0.6 | 16±11 | 4.7±0.8 | 0.6±0.5 |
| | T | 3.8±1.2 | 1.3±0.1 | 9±3.3 | 5.8±1.2 | 1.4±0.7 |
| $CD8^+$ | H | 28±2 | 14±13 | 38±4 | 29±16 | N/A |
| | I | 34±3 | 37±4 | 32±7 | 44±7 | 46±5 |
| | T | 29±4 | 35±3 | 24±4 | 40±2 | 42±7 |
| $CD8^+CD44^+$ | H | 28±1 | 18±4 | 60±5 | 60±13 | N/A |
| | I | 29±3 | 23±5 | 56±20 | 62±11 | 77±9 |
| | T | 20±2 | 31±5 | 53±14 | 63±3 | 71±16 |
| $CD8^+CD44^+ CXCR5^+$ | H | 1.6±1 | 1.6±0.9 | 0.5±0.1 | 6±3 | N/A |
| | I | 1±0.4 | 2.4±1.5 | 4±2 | 8.4±3.5 | 9.6±1.7 |
| | T | 1.5±0.8 | 5±2 | 2±3 | 7.6±2.3 | 9.6±2.8 |
| $CD8^+CD44^+CXCR5^+PD1^+$ | H | 0.6±0.2 | 0.02±0.01 | 1.7±1 | 0.2±0.3 | N/A |
| | I | 0.9±0.1 | 0.05±0.02 | 1.7±1.6 | 0.3±0.1 | 2.3±0.1 |
| | T | 0.5±0.2 | 0.07±0.03 | 0.9±0.4 | 0.3±0.2 | 2.8±1.2 |

**B**

| Lung cells | 2M | 4M | 17M | 19M |
|---|---|---|---|---|
| $CD4^+$ | 28±6 | 41±8 | 26±2 | 48±13 |
| $CD4^+CD44^+$ | 47±1 | 65±5 | 75±4 | 69±9 |
| $CD4^+CD44^+CXCR5^+$ | 3±1 | 4±3 | 3.6±1.7 | 5±2 |
| $CD4^+CD44^+CXCR5^+PD1^+$ | 6±3 | 1.1±1 | 2±1 | 0.8±0.6 |
| $CD8^+$ | 17±3 | 32±9 | 18±4 | 30±7 |
| $CD8^+CD44^+$ | 45±15 | 57±10 | 82±4 | 64±15 |
| $CD8^+CD44^+CXCR5^+$ | 0.9±0.3 | 3.6±0.9 | 6±3 | 4.3±1.7 |
| $CD8^+CD44^+CXCR5^+PD1^+$ | 4±2 | 0.16±0.1 | 2±1 | 0.14±0.1 |

**Supplementary Table S2:** Total number of sorted activated splenic $CD4^+$ T cells and their protein amount from healthy young (4 months) and old (19 months) C57BL/6 mice upon Mtb H37Rv infection and RIF -INH treatment. *This table is an extension of Fig. 3.*

| Mice Condition | Number of $CD4^+CD44^+$ cells sorted | Total protein amount (µg) |
|---|---|---|
| Healthy (Young) | 4,31,308 | 24.53 |
| | 3,23,580 | 13.96 |
| | 4,41,483 | 9.06 |
| | 4,35,189 | 20.38 |
| Mtb-infected (Young) | 5,82,869 | 15.47 |
| | 3,50,937 | 6.79 |
| | 3,00,415 | 6.42 |
| | 2,80,196 | 16.60 |
| | 1,63,427 | 7.17 |
| | 2,00,368 | 20.00 |
| RIF-INH treated (Young) | 2,47,515 | 13.58 |
| | 3,20,773 | 24.91 |
| | 6,49,437 | 18.11 |
| | 1,36,499 | 1.13 |
| | 5,18,395 | 6.79 |
| Healthy (Old) | 5,34,188 | 10.94 |
| | 3,61,425 | 15.47 |
| | 10,17,677 | 19.25 |
| | 6,63,651 | 13.21 |
| Mtb-infected (Old) | 4,55,972 | 8.30 |
| | 4,37,661 | 14.72 |
| | 4,28,506 | 12.45 |
| | 6,22,087 | 20.38 |
| | 6,84,220 | 10.94 |
| | 3,67,149 | 10.57 |
| RIF-INH treated (Old) | 7,38,297 | 6.79 |
| | 3,66,620 | 36.23 |
| | 1,99,573 | 36.60 |
| | 3,51,695 | 8.68 |

**Supplementary Table S3: List of identified proteins (n= 109) from splenic CD4$^{+}$CD44$^{+}$ T cells (TMT- Set 1) showing Mtb H37Rv infected young and old C57BL/6 mice with their respective healthy controls**. Y: young (4 months); O: old (19 months); I: Mtb infected; H: Healthy. *This table is an extension of Fig. 3.*

| Accession | Identified Protein(s) name | $log_2$fold change | | | | | | | |
|---|---|---|---|---|---|---|---|---|---|
| | | **YI1/YH2** | **YI2/YH2** | **YI3/YH2** | **YH1/YH2** | **OI1/OH2** | **OI2/OH2** | **OI3/OH2** | **OH1/OH2** |
| P68254 | 14-3-3 protein theta | 1.66 | 1.01 | 1.46 | -0.12 | 0.95 | 1.12 | 1.66 | 0.47 |
| P63101 | 14-3-3 protein zeta/delta | 2.12 | 1.53 | 2.35 | 0.86 | 2.32 | 1.17 | 2.58 | 0.70 |
| Q921R2 | 40S ribosomal protein S13 | -0.27 | 1.00 | 1.19 | 0.11 | -1.35 | -1.03 | 0.08 | 0.33 |
| Q9DCU9 | 4-hydroxy-2-oxoglutarate aldolase, mitochondrial | -1.45 | -0.98 | -2.35 | -0.75 | -2.04 | -1.97 | -2.13 | -2.09 |
| F6Z0X0 | 60S ribosomal protein L28 (Fragment) | -0.16 | 0.31 | 1.75 | 0.92 | 3.42 | 2.73 | 2.05 | 2.23 |
| Q9EST5 | Acidic leucine-rich nuclear phosphoprotein 32 family member B | 0.94 | 0.97 | 2.05 | 1.31 | 0.32 | 0.99 | 0.70 | -0.52 |
| Q9JM76 | Actin-related protein 2/3 complex subunit 3 | 1.86 | 2.12 | 2.31 | -0.11 | 1.30 | 1.85 | 1.73 | 1.26 |
| Q3UA72 | Actin-related protein 2/3 complex subunit 5 | 0.04 | -0.36 | 0.33 | -0.70 | 0.78 | 0.13 | 0.92 | 0.86 |
| Q9CPW4 | Actin-related protein 2/3 complex subunit 5 | 0.04 | -0.36 | 0.33 | -0.70 | 0.78 | 0.13 | 0.92 | 0.86 |
| A0A140LIG7 | Adaptor-related protein complex 2, alpha 1 subunit (Fragment) | -1.96 | -1.71 | -3.40 | -2.08 | -2.80 | -2.06 | -3.14 | -1.02 |
| P40124 | Adenylyl cyclase-associated protein 1 | 0.79 | 0.74 | -0.15 | -0.31 | -0.49 | -0.20 | 2.45 | 0.21 |
| P48962 | ADP/ATP translocase 1 | 1.60 | 0.89 | 0.77 | -0.23 | -1.22 | 1.49 | -0.22 | 0.49 |
| P84078 | ADP-ribosylation factor 1 | -0.72 | -1.43 | -2.10 | -1.86 | -1.19 | 0.60 | -1.15 | 0.32 |
| P07724 | Albumin | -3.24 | -1.89 | -4.74 | -3.08 | -0.57 | -2.00 | -2.07 | 1.12 |
| A0A5F8MPK3 | Ankyrin repeat and ubiquitin domain containing 1 | -0.03 | -1.30 | -2.06 | 0.23 | -1.53 | -1.16 | -3.56 | 0.66 |
| P17426 | AP-2 complex subunit alpha-1 | -1.96 | -1.71 | -3.40 | -2.08 | -2.80 | -2.06 | -3.14 | -1.02 |
| Q00623 | Apolipoprotein A-I | -2.97 | -1.41 | -6.18 | -4.63 | -1.61 | -1.28 | 1.08 | 2.50 |
| P56480 | ATP synthase subunit beta, mitochondrial | 0.30 | -0.30 | 0.04 | 0.76 | -0.30 | 0.14 | -0.46 | -0.43 |
| P58545 | BTB/POZ domain-containing protein 3 | 0.25 | 0.28 | -1.00 | 1.42 | -0.28 | -1.08 | -3.33 | -0.05 |
| P14211 | Calreticulin | 2.17 | 1.96 | 2.82 | 2.33 | 1.17 | 1.58 | 1.26 | 0.47 |
| Q9Z1Q5 | Chloride intracellular channel protein 1 | -0.15 | -1.05 | -1.87 | -0.71 | -1.20 | -0.36 | -0.60 | 0.35 |
| Q9CQI6 | Coactosin-like protein | 2.00 | 1.50 | 2.43 | 1.31 | 2.37 | 2.82 | 2.83 | 1.13 |

| | | | | | | | | | |
|---|---|---|---|---|---|---|---|---|---|
| P01029 | Complement C4-B | -1.79 | -1.02 | -2.01 | -0.05 | -2.34 | -1.99 | -2.26 | -1.14 |
| Q9QZQ8 | Core histone macro-H2A.1 | 0.32 | 0.74 | 0.85 | 0.68 | 1.16 | 0.71 | 1.81 | 0.77 |
| Q8CCK0 | Core histone macro-H2A.2 | 0.51 | 0.96 | 0.90 | 0.86 | 1.85 | -0.16 | 2.20 | 0.49 |
| G3UYK8 | Coronin | 1.95 | -0.02 | 0.52 | 0.14 | -1.26 | 1.92 | 2.62 | -0.28 |
| O89053 | Coronin-1A | 1.68 | -0.16 | -0.14 | -0.36 | -1.24 | 1.23 | 2.12 | -0.37 |
| P10126 | Elongation factor 1-alpha 1 | 1.24 | 0.67 | -0.03 | -1.32 | -0.51 | 0.79 | 2.72 | 0.07 |
| P62631 | Elongation factor 1-alpha 2 | 1.12 | 0.29 | -0.20 | -1.41 | -0.79 | 0.77 | 2.51 | -0.34 |
| Q8BGY2 | Eukaryotic translation initiation factor 5A-2 | 0.83 | 0.71 | 1.02 | -0.91 | 2.04 | 1.91 | 2.50 | 2.52 |
| P26040 | Ezrin | 0.75 | -0.39 | -0.67 | -1.72 | -0.81 | 0.42 | 1.38 | -0.50 |
| P16045 | Galectin-1 | 3.21 | 2.53 | 2.83 | 0.62 | 0.85 | 0.27 | 0.73 | -1.11 |
| Q9ERL7 | Glia maturation factor gamma | 0.62 | -0.53 | -0.37 | -0.06 | 1.11 | 1.09 | -1.25 | 0.52 |
| Q64467 | Glyceraldehyde-3-phosphate dehydrogenase, testis-specific | 1.52 | 1.20 | 1.90 | -0.07 | 2.64 | 2.73 | 3.81 | 1.56 |
| P62827 | GTP-binding nuclear protein Ran | 1.70 | 0.75 | 1.05 | -0.49 | 1.66 | 1.08 | 1.76 | 0.31 |
| Q9D2U9 | H2B.U histone 2 | 1.43 | 1.72 | 2.04 | 1.38 | 3.09 | 2.96 | 3.25 | 2.02 |
| P11499 | Heat shock protein HSP 90-beta | 1.52 | 0.31 | 0.59 | -0.43 | -1.68 | 0.64 | 1.22 | -0.44 |
| P01942 | Hemoglobin subunit alpha | -0.61 | -0.84 | -3.25 | -3.36 | -2.11 | -0.27 | 0.80 | 2.36 |
| P02089 | Hemoglobin subunit beta-2 | -0.41 | -1.61 | -2.36 | -2.36 | -1.53 | 0.04 | -0.26 | 1.65 |
| P06467 | Hemoglobin subunit zeta | -0.51 | 0.28 | -3.73 | -2.75 | -3.19 | -2.34 | -0.59 | 2.53 |
| A0A0N4SUM2 | Heterogeneous nuclear ribonucleoprotein A2/B1 (Fragment) | -0.44 | 1.10 | -1.04 | -0.84 | -0.17 | -1.87 | 0.71 | -0.66 |
| Q60668 | Heterogeneous nuclear ribonucleoprotein D0 | 1.18 | 0.84 | 1.46 | 0.71 | 3.20 | 2.60 | 4.06 | 2.05 |
| H3BLL4 | Heterogeneous nuclear ribonucleoprotein K | 0.91 | 0.61 | 0.12 | -0.78 | -0.09 | 0.76 | 2.32 | 0.58 |
| Q8VEK3 | Heterogeneous nuclear ribonucleoprotein U | 0.30 | -0.12 | -0.80 | -0.27 | -2.58 | -1.64 | 1.66 | -0.61 |
| O88569 | Heterogeneous nuclear ribonucleoproteins A2/B1 | 0.41 | 3.24 | -0.34 | -1.07 | -0.38 | -1.31 | 3.30 | 0.82 |
| P30681 | High mobility group protein B2 | 2.17 | 1.67 | 3.63 | 1.34 | 2.29 | 3.44 | 1.87 | 0.16 |
| Q9JHD1 | Histone acetyltransferase KAT2B | -0.82 | 5.28 | 3.75 | -0.37 | -9.30 | -7.78 | -4.06 | -1.77 |
| P43275 | Histone H1.1 | 0.68 | 0.54 | 1.83 | 0.96 | 4.63 | 3.44 | 3.65 | 3.18 |
| P15864 | Histone H1.2 | -0.10 | 1.78 | 3.26 | 0.38 | 2.67 | 1.43 | 4.77 | 1.72 |
| P43274 | Histone H1.4 | -0.76 | 0.30 | -0.19 | -0.45 | 4.92 | 4.04 | 4.52 | 3.79 |
| P43276 | Histone H1.5 | -0.97 | 1.11 | -1.31 | -1.50 | 4.62 | 3.14 | 5.11 | 3.57 |

| | | | | | | | | | |
|---|---|---|---|---|---|---|---|---|---|
| Q8CGP5 | Histone H2A type 1-F | 1.08 | 1.13 | 2.34 | 0.84 | 2.67 | 2.19 | 2.39 | 0.81 |
| Q8CGP7 | Histone H2A type 1-K | 1.08 | 1.13 | 2.34 | 0.84 | 2.67 | 2.19 | 2.39 | 0.81 |
| Q6GSS7 | Histone H2A type 2-A | 1.35 | 0.20 | 2.91 | 2.16 | 2.38 | 2.02 | 1.48 | -0.74 |
| Q64523 | Histone H2A type 2-C | -0.03 | -0.35 | 0.81 | -0.25 | 0.83 | 0.48 | 0.97 | 0.11 |
| P0C0S6 | Histone H2A.Z | 0.85 | 0.33 | 1.64 | 0.58 | -0.11 | 0.51 | -1.00 | -1.67 |
| Q6ZWY9 | Histone H2B type 1-C/E/G | 1.08 | 1.40 | 1.69 | 0.51 | 2.66 | 2.54 | 2.83 | 1.44 |
| Q8CGP0 | Histone H2B type 3-B | 1.43 | 1.72 | 2.04 | 1.38 | 3.09 | 2.96 | 3.25 | 2.02 |
| P62806 | Histone H4 | 2.17 | 2.68 | 2.25 | 1.50 | 1.25 | 1.82 | 1.05 | 1.11 |
| A6X935 | Inter alpha-trypsin inhibitor, heavy chain 4 | -0.70 | -0.54 | -2.38 | 0.63 | -1.01 | -1.31 | -2.37 | -0.34 |
| A0A2I3BRQ3 | Inter-alpha trypsin inhibitor, heavy chain 3 | -0.83 | -0.24 | -1.44 | 0.30 | -1.83 | -1.40 | -1.25 | -1.00 |
| Q61704 | Inter-alpha-trypsin inhibitor heavy chain H3 | -0.94 | -0.39 | -1.62 | 0.38 | -1.68 | -1.48 | -1.32 | -0.75 |
| A0A1B0GSL7 | Lactate dehydrogenase A (Fragment) | 1.62 | 0.78 | 0.83 | -1.14 | -0.25 | 0.92 | 0.94 | -0.23 |
| P08071 | Lactotransferrin | 1.80 | -1.20 | 1.62 | 2.06 | -0.36 | 2.40 | -2.68 | -1.57 |
| Q61029 | Lamina-associated polypeptide 2, isoforms beta/delta/epsilon/gamma | -1.26 | -1.06 | 5.25 | -0.54 | -4.81 | -4.59 | -1.97 | -3.82 |
| P61358 | Large ribosomal subunit protein eL27 | 2.33 | 2.35 | 3.57 | -1.40 | 1.21 | 1.27 | 1.45 | 0.91 |
| P41105 | Large ribosomal subunit protein eL28 | -0.16 | 0.31 | 1.75 | 0.92 | 3.42 | 2.73 | 2.05 | 2.23 |
| O55142 | Large ribosomal subunit protein eL33 | 1.00 | 1.83 | 0.37 | 0.18 | 2.55 | 1.03 | 2.34 | 1.45 |
| P47911 | Large ribosomal subunit protein eL6 | 0.75 | 0.78 | 0.92 | -0.14 | 1.45 | 1.06 | 1.67 | 0.13 |
| D3YZQ9 | L-lactate dehydrogenase (Fragment) | 1.14 | 1.35 | 0.46 | -0.49 | 0.13 | 0.49 | 0.96 | 0.50 |
| A0A0N4SVV8 | L-lactate dehydrogenase (Fragment) | 0.58 | 1.80 | 0.06 | 0.00 | 0.46 | 0.01 | 0.97 | 1.04 |
| A0A1B0GQX5 | L-lactate dehydrogenase | 1.14 | 1.35 | 0.46 | -0.49 | 0.13 | 0.49 | 0.96 | 0.50 |
| A0A6I8MX27 | L-lactate dehydrogenase | 0.58 | 1.80 | 0.06 | 0.00 | 0.46 | 0.01 | 0.97 | 1.04 |
| D3Z311 | Lymphocyte cytosolic protein 1 (Fragment) | 1.00 | -0.04 | 0.11 | -2.06 | -0.42 | 1.47 | 4.36 | 1.08 |
| A2ANY6 | Midasin | -0.92 | -0.68 | -2.32 | -1.98 | 0.58 | 0.06 | 1.12 | 2.28 |
| P26041 | Moesin | 0.92 | 0.54 | 0.01 | -0.27 | -0.18 | 0.63 | 1.74 | -0.38 |
| Q60605 | Myosin light polypeptide 6 | 0.98 | 0.48 | 1.86 | 0.79 | 0.81 | 0.56 | 0.44 | -0.26 |
| Q61937 | Nucleophosmin | 1.98 | 2.10 | 2.10 | 1.34 | 2.93 | 2.89 | 2.76 | 1.06 |
| Q5NC80 | Nucleoside diphosphate kinase (Fragment) | 0.09 | 0.11 | -1.03 | -1.05 | -0.34 | 0.80 | -0.04 | -0.11 |
| Q60887 | Olfactory receptor 10N1 | 2.66 | 2.87 | 2.82 | 1.51 | 1.97 | 3.03 | 1.89 | 1.74 |
| P17742 | Peptidyl-prolyl cis-trans isomerase A | -0.75 | -0.13 | -1.50 | 0.16 | -0.93 | -1.23 | 0.25 | -1.02 |

| | | | | | | | | | |
|---|---|---|---|---|---|---|---|---|---|
| Q61233 | Plastin-2 | 1.00 | -0.04 | 0.11 | -2.06 | -0.42 | 1.47 | 4.36 | 1.08 |
| Q91Z31 | Polypyrimidine tract-binding protein 2 | -0.13 | -0.38 | 0.17 | 0.32 | -1.12 | -0.08 | -0.56 | -0.42 |
| P62962 | Profilin-1 | 0.82 | 2.07 | 2.77 | 1.64 | 0.68 | 1.90 | -0.10 | 0.94 |
| D3Z7C6 | Prostaglandin E synthase 3 | 0.22 | -0.50 | -0.23 | -0.20 | 1.18 | 0.93 | 1.34 | 0.10 |
| P28063 | Proteasome subunit beta type-8 | 2.01 | 2.25 | 3.74 | 2.41 | 1.33 | 1.75 | 2.28 | 0.86 |
| P07091 | Protein S100-A4 | 2.46 | 2.31 | 2.02 | -0.74 | -3.14 | -2.93 | -0.49 | -3.41 |
| Q64G17 | Putative acidic leucine-rich nuclear phosphoprotein 32 family member C | 0.94 | 0.97 | 2.05 | 1.31 | 0.32 | 0.99 | 0.70 | -0.52 |
| A0A5F8MPB9 | Radixin | 0.89 | 0.23 | -0.31 | -1.14 | -0.33 | 0.85 | 1.81 | 0.04 |
| Q7TSG6 | Radixin | 0.75 | -0.39 | -0.67 | -1.72 | -0.81 | 0.42 | 1.38 | -0.50 |
| P26043 | Radixin | 0.89 | 0.23 | -0.31 | -1.14 | -0.33 | 0.85 | 1.81 | 0.04 |
| Q99PT1 | Rho GDP-dissociation inhibitor 1 | 1.38 | 0.67 | 0.86 | 0.84 | 2.01 | 1.96 | 0.82 | 1.60 |
| Q61599 | Rho GDP-dissociation inhibitor 2 | 0.60 | 0.73 | 0.16 | -1.15 | 0.80 | 0.36 | 2.36 | 1.20 |
| Q921I1 | Serotransferrin | -1.73 | -1.43 | -3.51 | -0.72 | -2.23 | -3.38 | -4.54 | -1.28 |
| P97351 | Small ribosomal subunit protein eS1 | 0.79 | 0.54 | 1.23 | 0.01 | 1.30 | 1.63 | 1.78 | 0.90 |
| P68040 | Small ribosomal subunit protein RACK1 | 0.93 | 0.20 | -0.20 | 1.49 | -1.44 | -0.42 | 0.31 | 0.79 |
| P62270 | Small ribosomal subunit protein uS13 | 0.06 | -0.16 | 0.79 | -0.74 | 0.39 | -1.01 | -0.07 | 0.26 |
| P62301 | Small ribosomal subunit protein uS15 | -0.27 | 1.00 | 1.19 | 0.11 | -1.35 | -1.03 | 0.08 | 0.33 |
| Q6ZWN5 | Small ribosomal subunit protein uS4 | 2.33 | 1.65 | 3.03 | 1.76 | 1.31 | 1.47 | 0.79 | -0.29 |
| P14131 | Small ribosomal subunit protein uS9 | 2.26 | 1.66 | 2.58 | 1.08 | 1.32 | 1.46 | 1.31 | 0.34 |
| Q8C9H6 | Striatin-interacting proteins 2 | -1.45 | -0.98 | -2.35 | -0.75 | -2.04 | -1.97 | -2.13 | -2.09 |
| P08228 | Superoxide dismutase [Cu-Zn] | 2.86 | 2.05 | 2.74 | 1.35 | -0.53 | 1.48 | 1.20 | -0.97 |
| O08583 | THO complex subunit 4 | 2.41 | 0.45 | 2.70 | 1.66 | 0.12 | 1.60 | -1.54 | -1.48 |
| P20065 | Thymosin beta-4 | -0.27 | 0.88 | -2.37 | -0.93 | 1.57 | 0.74 | 2.95 | 0.62 |
| B7ZNL3 | Tpm1 protein | 0.93 | 0.32 | -0.12 | 0.06 | 0.54 | 2.03 | 0.95 | -0.23 |
| Q9WVA4 | Transgelin-2 | -0.07 | -0.61 | -0.96 | 0.03 | -1.77 | -1.04 | -2.12 | -0.89 |
| Q9JKK7 | Tropomodulin-2 | 0.65 | 0.63 | -2.90 | -0.08 | -2.78 | -1.25 | -3.31 | 0.80 |
| Q8BSH3 | Tropomyosin 1, alpha | 0.93 | 0.32 | -0.12 | 0.06 | 0.54 | 2.03 | 0.95 | -0.23 |
| Q9EQH3 | Vacuolar protein sorting-associated protein 35 | -1.09 | 1.47 | -2.61 | -1.38 | -2.28 | -1.66 | -0.99 | -1.45 |

**Supplementary Table S4: List of identified proteins (n=41) from splenic CD4$^{+}$CD44$^{+}$ T cells (TMT- Set 2) showing RIF-INH treated young and old C57BL/6 mice with Mtb H37Rv infected controls**. Y: young (4 months); O: old (19 months); T: RIF-INH treated; I: Mtb infected. *This table is an extension of Fig. 3.*

| Accession | Identified Protein(s) name | $log_2$fold change | | | | | | | |
|---|---|---|---|---|---|---|---|---|---|
| | | **YT1/YI1** | **YT2/YI1** | **YT3/YI1** | **YI2/YI1** | **OT1/OI1** | **OT2/OI1** | **OT3/OI1** | **OI2/OI1** |
| P50247 | Adenosylhomocysteinase | -1.05 | 0.04 | -4.26 | -1.06 | -1.85 | 0.55 | 1.47 | -0.50 |
| P32020 | Sterol carrier protein 2 | -0.88 | -1.28 | -3.18 | -0.32 | -1.27 | -0.80 | 0.37 | -1.20 |
| A0A0G2JGN4 | Small nuclear ribonucleoprotein B | -1.17 | -1.59 | -2.98 | -0.86 | -1.27 | -1.28 | 1.24 | 0.24 |
| S4R1W1 | Glyceraldehyde-3-phosphate dehydrogenase | -2.30 | -2.27 | -1.51 | -1.88 | -0.41 | -0.22 | -1.00 | 0.50 |
| O08583 | THO complex subunit 4 | -0.53 | -0.80 | -0.01 | 0.54 | -3.02 | -3.39 | -0.11 | -0.55 |
| P62806 | Histone H4 | -1.29 | -3.34 | -3.27 | -0.02 | -1.88 | -3.44 | -0.43 | -1.12 |
| Q64524 | Histone H2B type 2-E | -0.70 | -1.60 | -2.52 | 0.07 | -2.39 | -4.07 | 0.24 | 0.10 |
| F8WI35 | Histone H3 | -2.11 | -2.57 | -2.79 | -0.18 | -1.91 | -3.82 | -1.45 | 0.48 |
| E0CZ27 | Histone H3 (Fragment) | -2.18 | -2.62 | -2.80 | -0.19 | -1.86 | -3.80 | -1.44 | 0.53 |
| Q64525 | Histone H2B type 2-B | -1.24 | -2.24 | -2.69 | -0.16 | -1.75 | -3.49 | -1.25 | 0.62 |
| Q8CGP2 | Histone H2B type 1-P | -0.96 | -1.63 | -2.41 | 0.05 | -2.00 | -3.58 | -0.13 | 0.24 |
| Q9D2U9 | H2B.U histone 2 | -0.94 | -1.79 | -2.60 | 0.01 | -2.03 | -3.88 | 0.00 | 0.20 |
| E0CYR7 | H3.3 histone A (Fragment) | -2.11 | -2.65 | -2.87 | -0.20 | -1.71 | -3.76 | -1.34 | 0.57 |
| Q8CGP0 | Histone H2B type 3-B | -0.94 | -1.79 | -2.60 | 0.01 | -2.03 | -3.88 | 0.00 | 0.20 |
| P70696 | Histone H2B type 1-A | -0.96 | -1.65 | -2.36 | 0.04 | -1.95 | -3.53 | -0.17 | 0.25 |
| A0A8I4SYN6 | Histone H3 | -2.00 | -2.43 | -2.66 | -0.15 | -1.51 | -3.78 | -1.50 | 0.36 |
| P27661 | Histone H2AX | -2.19 | -2.60 | -2.65 | -0.37 | -1.17 | -3.93 | -0.60 | 0.91 |
| P07309 | Transthyretin | 0.39 | 0.03 | 0.55 | -0.69 | 1.41 | 1.74 | 0.88 | -0.61 |
| V9GX81 | Maestro heat-like repeat family member 6 | -0.35 | -0.12 | -0.64 | 0.21 | 0.04 | -0.17 | -1.56 | -0.12 |
| P15864 | Histone H1.2 | -0.99 | -0.48 | -1.84 | 0.27 | -2.37 | -4.25 | -3.62 | 0.06 |
| Q07133 | Histone H1t | -1.41 | -0.61 | -2.62 | 0.14 | -2.12 | -5.30 | -4.39 | 0.09 |
| P43275 | Histone H1.1 | -0.49 | -0.29 | -1.86 | 0.56 | -1.61 | -3.81 | -4.33 | 0.12 |
| G3UWL7 | Histone H2A | -2.03 | -2.53 | -2.37 | -0.67 | 0.52 | -4.01 | -0.79 | 1.19 |

| | | | | | | | | | |
|---|---|---|---|---|---|---|---|---|---|
| B2RY04 | Dedicator of cytokinesis protein 5 | -0.06 | -0.51 | 0.80 | 0.31 | 2.06 | 2.02 | | -0.65 |
| A8DUK4 | Beta-globin | 6.13 | 5.47 | 3.53 | -0.15 | -1.76 | 1.11 | 2.33 | -1.13 |
| A0A0U1RQ96 | Actin, gamma 2, smooth muscle, enteric (Fragment) | -4.50 | -4.81 | -4.38 | -2.37 | -4.75 | -3.76 | -3.22 | 0.85 |
| Q8BFZ3 | Beta-actin-like protein 2 | -3.66 | -3.91 | -4.29 | -2.11 | -3.98 | -3.68 | -2.62 | 0.79 |
| E9Q5F4 | Actin, beta (Fragment) | -3.91 | -3.75 | -1.03 | -1.72 | -2.24 | -3.14 | -2.50 | 0.38 |
| E9Q1F2 | Actin, beta | -4.02 | -3.82 | -0.94 | -1.72 | -2.21 | -3.12 | -2.55 | 0.36 |
| P63268 | Actin, gamma-enteric smooth muscle | -3.67 | -3.60 | -0.67 | -1.64 | -2.18 | -3.22 | -2.43 | 0.21 |
| A0A1D5RM20 | Actin alpha 1, skeletal muscle (Fragment) | -3.56 | -3.46 | -0.41 | -1.55 | -1.96 | -3.15 | -2.33 | 0.10 |
| F6WX90 | Actin, alpha, cardiac muscle 1 (Fragment) | -3.67 | -3.53 | -0.28 | -1.55 | -1.90 | -3.13 | -2.39 | 0.06 |
| A0A338P692 | Alpha-2-HS-glycoprotein (Fragment) | -0.79 | -0.31 | -0.47 | 0.10 | 0.78 | 1.85 | -3.15 | -0.10 |
| Q6NS59 | Protein FAM135A | 0.34 | 0.44 | 0.37 | 0.42 | 0.90 | 1.98 | -4.33 | -0.17 |
| Q3T052 | Inter-alpha-trypsin inhibitor heavy chain H4 | -0.57 | -0.34 | -0.06 | 0.09 | 1.17 | 0.37 | -3.07 | 0.25 |
| P06467 | Hemoglobin subunit zeta | -0.57 | -2.06 | -3.20 | -0.70 | 0.75 | 3.25 | -0.33 | -0.41 |
| F7CJN9 | Transferrin (Fragment) | -0.28 | 0.59 | 0.74 | 0.09 | 2.19 | 2.57 | -2.00 | -0.21 |
| Q921I1 | Serotransferrin | 0.40 | 0.87 | 0.52 | 0.09 | 1.96 | 1.85 | -0.36 | -0.45 |
| P57016 | Ladinin-1 | 0.74 | 0.77 | 0.81 | -0.54 | 2.30 | 1.96 | 1.31 | -1.21 |
| P05202 | Aspartate aminotransferase, mitochondrial | -0.63 | 0.50 | 1.24 | 0.58 | 3.23 | 2.87 | 3.35 | -0.24 |
| D3YYR8 | Transferrin (Fragment) | 2.24 | 1.89 | -0.89 | 0.07 | 1.50 | -0.47 | 1.17 | -0.97 |
